# Nanoscale surface topography reduces focal adhesions and cell stiffness by enhancing integrin endocytosis

**DOI:** 10.1101/2021.06.18.448920

**Authors:** Xiao Li, Lasse H. Klausen, Wei Zhang, Zeinab Jahed, Ching-Ting Tsai, Thomas L. Li, Bianxiao Cui

## Abstract

Both substrate stiffness and surface topography regulate cell behavior through mechanotransduction signaling pathways. Such intertwined effects suggest that engineered surface topographies might substitute or cancel the effects of substrate stiffness in biomedical applications. However, the mechanisms by which cells recognize topographical features are not fully understood. Here we demonstrate that the presence of nanotopography drastically alters cell behavior such that neurons and stem cells cultured on rigid glass substrates behave as if they were on soft hydrogels. We further show that rigid nanotopography resembles the effect of soft hydrogels in reducing cell stiffness and membrane tension as measured by atomic force microscopy. Finally, we demonstrate that nanotopography reduces focal adhesions and cell stiffness by enhancing the endocytosis and the subsequent removal of integrin receptors. This mechanistic understanding will support the rational design of nanotopography that directs cells on rigid materials to behave as if they were on soft ones.

**TOC graphic:** 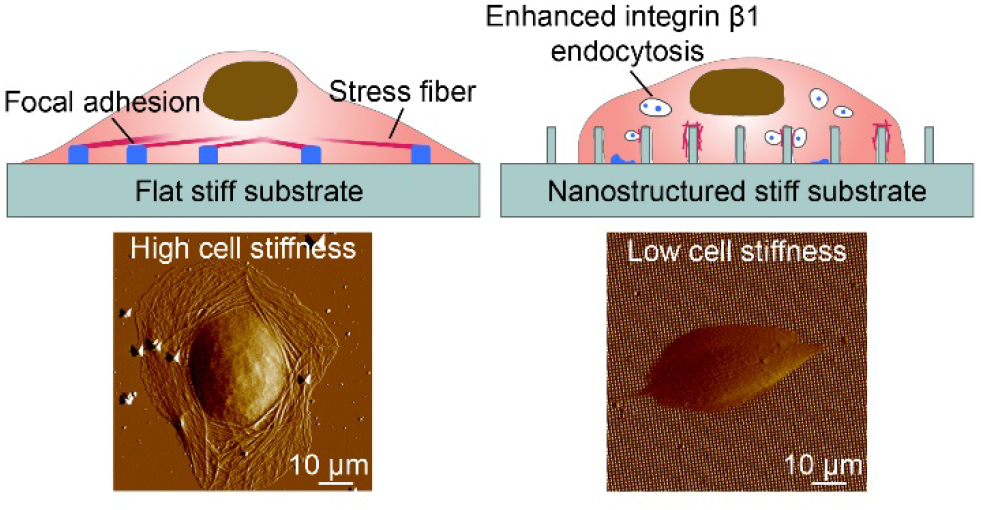

---

The stiffness of substrates has been shown to affect a wide range of cell behaviors such as stem cell differentiation,^1–3^ aging,^4^ and cancer cell invasion.^5^ In response to substrate stiffness, cells remodel their actin cytoskeleton and adapt their own stiffness and membrane tension.^6, 7^ Underlying these changes is the mechanotransduction machinery that exerts cellular traction forces onto extracellular materials through integrin receptors, whose activities are force-dependent.^8^ Rigid substrates can withstand high traction forces for enhanced integrin activation, which results in increased signaling cascades in mechanotransduction. From the application point of view, most human tissues exhibit low stiffness (e.g., Young’s modulus ~1 kPa for brain tissues^9^) while many biomedical materials are high stiffness (~120 GPa for titanium implants). This stiffness disparity increases foreign body responses^10–12^ and causes cell behaviors on rigid materials to differ from those in native soft microenvironments.^13, 14^ To meet this challenge, extensive efforts have been dedicated to developing soft materials, such as hydrogels, for interfacing with cells and tissues.^15^ However, soft materials do not meet all the demands of biodevices, such as implants that ought to provide mechanical support^16, 17^ and bioelectronic devices that rely on noble metals.^18^

Alongside stiffness, surface topography is another important physical property of extracellular materials. It has been shown that nanoscale surface topography influences cell behaviors, such as adhesion,^19^ alignment,^20–22^ migration,^23^ and differentiation.^24–26^ Some studies revealed that nanoscale surface topography reduces actin stress fibers and focal adhesions,^27–29^ which are key components of the mechanotransduction machinery. Other studies reported that nanotopography affects the activity of yes-associated protein (YAP), a nuclear regulator involved in mechanotransduction.^30, 31^ The documented effects of stiffness and nanotopography on mechanotransduction suggest that it may be possible to engineer surface topography to offset or reduce stiffness-induced effects on cells. However, surface topography is defined by a high-dimensional space of features, such as domain size, shape, height, steepness, and spacing. The molecular mechanisms underlying how cells recognize surface topography to modulate mechanotransduction is not well understood. This hinders the rational design of surface topography.

In this work, we focus on the effect of nanotopography, and we compare, from behavioral to molecular levels, the cellular responses separately induced by stiffness and nanotopography. We demonstrate that surface nanotopography can substantially modulate cellular responses to rigid glass substrates such that they are similar to responses to soft hydrogels. Furthermore, we reveal that a key mechanism of topography-sensing involves the endocytosis of integrin receptors, which is significantly enhanced by nanotopography-induced membrane deformation.

## Results and Discussion

### Rigid nanopillars accelerate neurite outgrowth in ways similar to soft hydrogels

We first examined how nanotopography and substrate stiffness differentially modulate the development of embryonic neurons. Quartz has a Young’s modulus around 72 GPa, whereas the polyacrylamide hydrogels used in comparison have a Young’s modulus of 1.04 kPa (named 1 kPa hydrogel). Well-defined nanotopography composed of quartz nanopillars were fabricated by electron beam lithography and reactive ion etching (**Figure 1a**, and **Figure S1**).^32^ Unlike vertical posts made of elastic materials,^2, 33^ these quartz nanopillars are rigid and not bendable.^34^ We began with nanopillars similar to those used in earlier studies (200 nm diameter, 1 μm pitch, and, 3 μm height). These nanopillars induced membrane wrapping as illustrated by the CellMask fluorescence imaging (**Figure S2**). By comparing cells on nanopillar areas and cells on flat areas of the same culture, we excluded variabilities associated with surface treatment and culture conditions.

**Figure 1.**
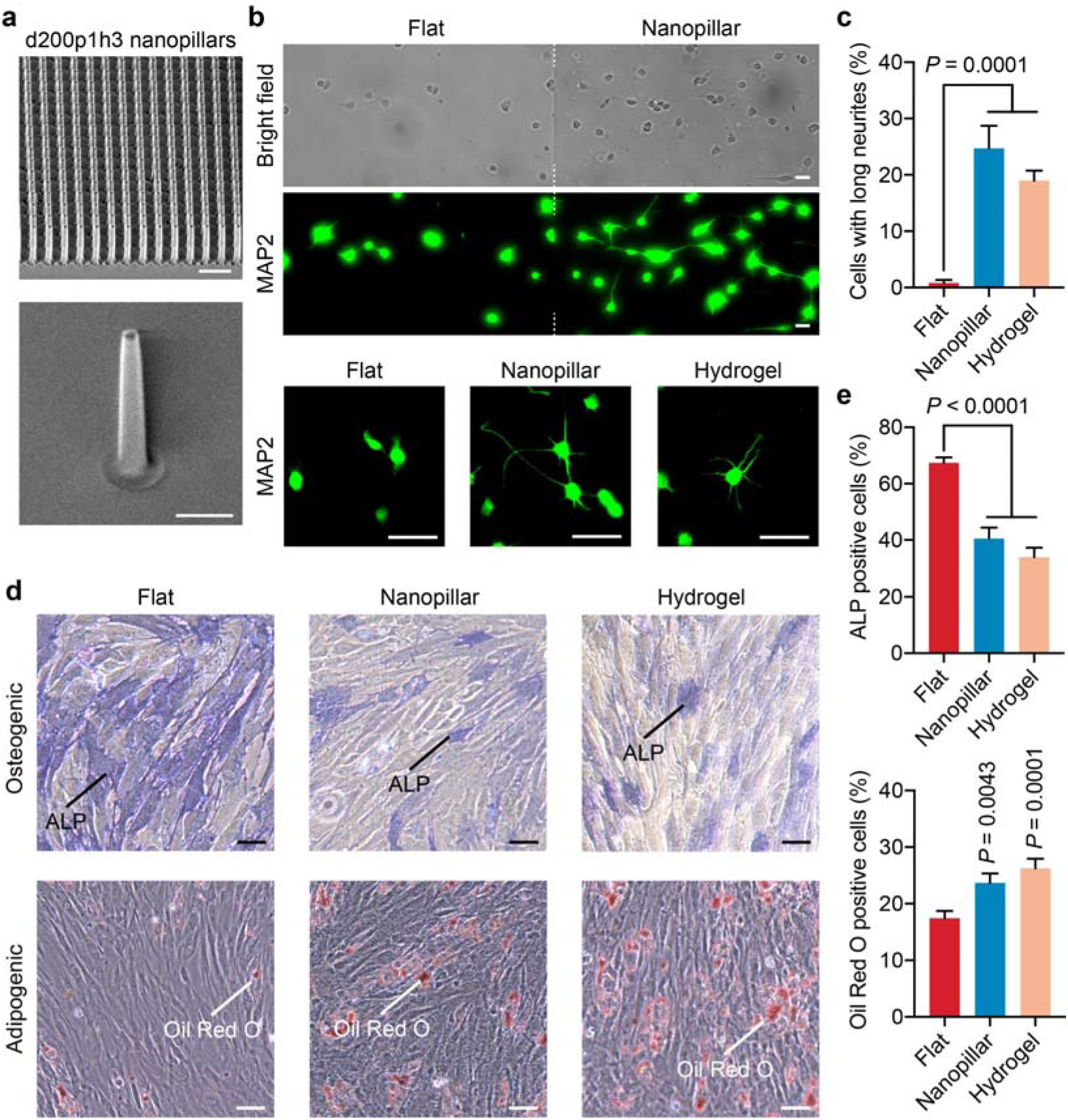
Quartz nanotopography induces similar cell behavior as soft hydrogels do. (a) Scanning electron microscopy (SEM) images of quartz nanopillars. The upper image shows an array of d200p1h3 nanopillars, and the lower image shows a single nanopillar. (b) Bright field and corresponding anti-MAP2 fluorescence images of E18 rat hippocampal neurons after 20 hr of culture on flat quartz (flat), nanopillar quartz (nanopillar), and 1 kPa hydrogel (hydrogel) surfaces. Upper images show neurons on a quartz substrate that contains both flat areas and nanopillar areas. White dash lines indicate the border between the flat and nanopillar areas. Lower images show neurons cultured on three different surfaces. (c) Quantified percentages of hippocampal neurons with at least one neurite longer than 35 μm (n = 193, 297, and 178 cells in flat, nanopillar, and hydrogel groups, respectively; mean ± s.e.m.). (d) Representative images of ALP staining and Oil Red O staining of hMSCs for osteogenic differentiation and adipogenic differentiation, respectively, on three different surfaces. (e) Percentage of hMSCs with positive ALP staining and positive Oil Red O staining, respectively (for ALP staining, n = 3856, 953, and 1353 cells in flat, nanopillar, and hydrogel groups, respectively; for Oil Red O staining, n = 5152, 5069, and 3819 cells in flat, nanopillar, and hydrogel groups, respectively; mean ± s.e.m.). *P* value in comparison to flat surface was determined by unpaired two-tailed t test. Scale bars, 2 μm (a, upper), 1 μm (a, lower), 20 μm (b), 50 μm (d).

Neurons are one of the softest cell types in the human body, and substrate stiffness has been shown to affect neuronal development.^14, 35, 36^ We measured neurite outgrowth in response to three types of surfaces: rigid flat quartz, rigid nanopillar quartz, and soft flat hydrogel surfaces. 20 hr after plating, embryonic E18 hippocampal neurons on flat quartz surfaces were mostly round with short extensions, consistent with previous studies using flat rigid substrates.^35^ In contrast, a significant fraction of neurons on nanopillar surfaces had long neurites in the same culture (**Figure 1b, S3**). Hippocampal neurons cultured on hydrogels also grew out long neurites after 20 hr, agreeing with previous reports using soft substrates.^14, 36^ Statistical analysis shows that the percentage of neurons having at least one neurite longer than 35 μm is significantly higher on nanopillars (24.7%) and 1 kPa hydrogels (19.0%) than on flat quartz (0.8%) (**Figure 1c**). These results demonstrate that nanotopography accelerates neurite outgrowth. We found that nanopillars of different dimensions also accelerated neurite outgrowth (**Figure S4**). A systematic study of how nanopillar dimensions affect cell behavior is presented in a later section.

### Rigid nanopillars bias stem cell differentiation in ways similar to soft hydrogels

We then examined how nanotopography and stiffness differentially modulate stem cell differentiation. Human mesenchymal stem cells (hMSCs) can differentiate into osteoblasts or adipocytes. Previous studies have separately shown that nanotopography and stiffness affect the differentiation efficiency of hMSCs into osteogenic or adipogenic lineages.^1, 29, 37, 38^ In particular, microscale topographic regulation of stem cell differentiation has been well established.^39, 40^ Here we compared hMSC differentiation in response to rigid flat quartz, rigid nanopillar quartz, and soft flat hydrogel surfaces. We first differentiated hMSCs toward the osteogenic lineage. 14 days after induction of differentiation, hMSCs were stained for alkaline phosphatase (ALP). As shown in **Figure 1d**, there was much less ALP staining on nanopillar and soft hydrogel surfaces compared to flat quartz surfaces. Quantitative analysis shows a dramatic reduction of ALP-positive cells from 67.3% on flat quartz to 40.5% on nanopillar quartz surfaces, which is close to 33.9% on 1 kPa hydrogel surfaces (**Figure 1e**).

We also examined the differentiation of hMSCs toward the softer adipogenic lineage. Six days after the differentiation induction, cells were stained with Oil Red O. We found that more cells were committed to the adipogenic lineage (positively stained in red) on nanopillars and hydrogels than on flat quartz surfaces (**Figure 1d**). Quantitative analysis confirms that hMSCs were differentiated into adipocytes with higher efficiency on nanopillars (23.7%) and hydrogels (26.3%) than on flat quartz surfaces (17.5%) (**Figure 1e)**. These results regarding neurite outgrowth and hMSC differentiation demonstrate the possibility of engineering surface topography such that cells on rigid quartz substrates behave like those on soft hydrogels.

### Rigid nanopillars reduce cell stiffness and membrane tension in ways similar to soft hydrogels

It is known that mammalian cells adjust their own mechanics according to the stiffness of extracellular materials.^6, 7, 41^ We used AFM to examine how nanotopography and substrate stiffness differentially affected cell stiffness and membrane tension. Contact-mode AFM shows that cells on flat quartz had visible cytoskeletal fibers surrounding the nucleus, but cells on nanopillars exhibited a drastically smaller shape with no discernable stress fibers (**Figure 2a**). For cell stiffness measurement, we used a large spherical probe and applied small indentations (< 500 nm) to ensure that a representative elasticity of the entire cell was obtained, where the elasticity of the cell cortex made the major contribution.^42^ Cell stiffness was extracted by fitting the Hertz indentation model to force–separation curves obtained from each cell **(Figures 2b, c, Figures S5a, b**). We found that cells on quartz nanopillars were significantly softer than cells on flat quartz with a 46% reduction of Young’s modulus and were comparable in stiffness to cells on 1 kPa hydrogels (**Figure 2d**).

**Figure 2.**
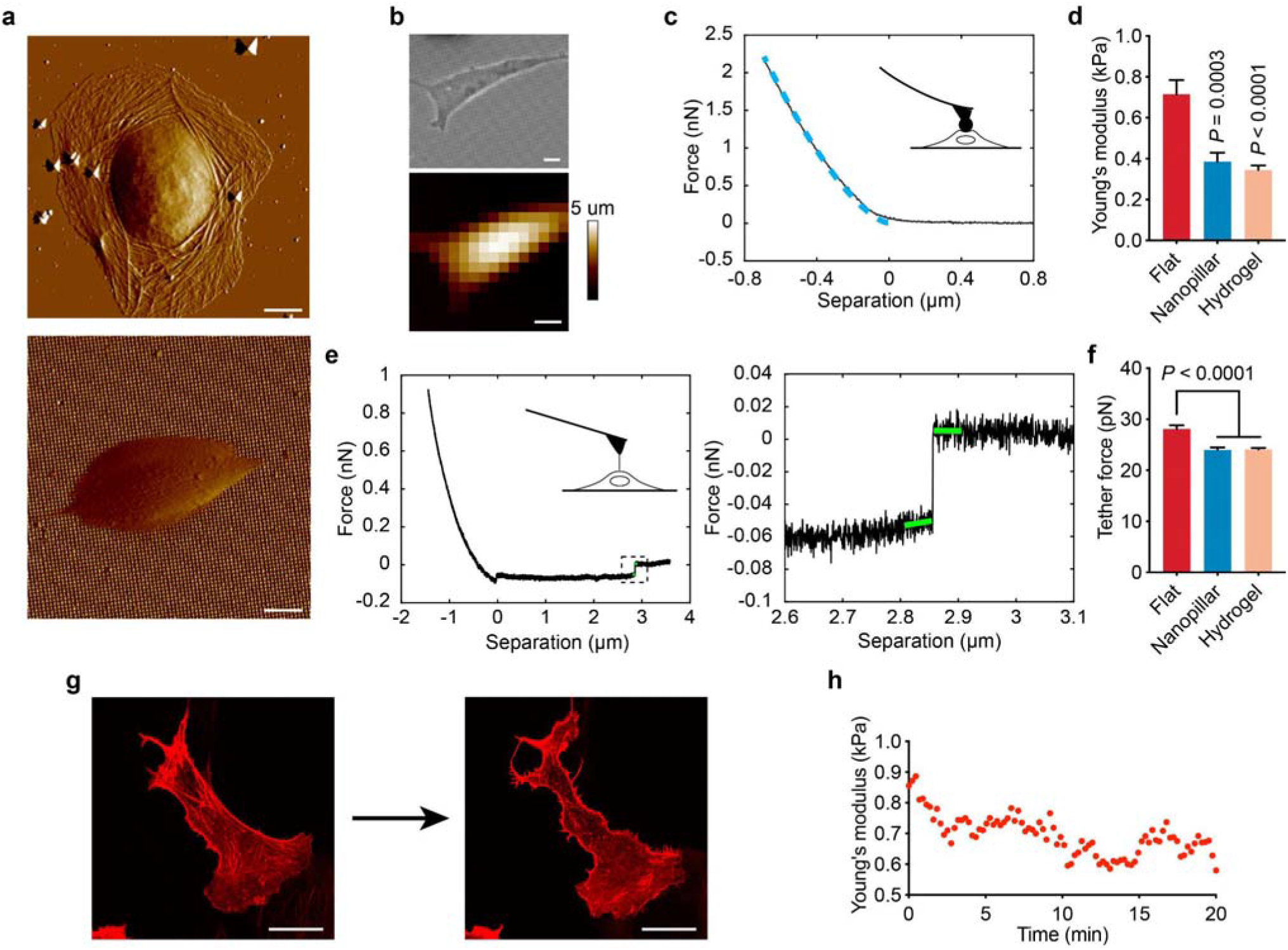
Quartz nanotopography reduces cell stiffness and membrane tension in a manner similar to soft hydrogels. (a) Contact-mode AFM deflection images of U2OS cells show different cell shapes on flat quartz (upper) and d200p1h3 nanopillar quartz (lower) surfaces. (b) Optical image (upper) and AFM-obtained topography map (lower) of a U2OS cell on d200p1h3 nanopillar surface. (c) A representative AFM force-separation curve obtained by AFM indentation. The blue dashed line indicates the Hertz fit used to calculate Young’s modulus. (d) Quantitative analysis of the Young’s modulus of cells obtained by AFM indentation on flat quartz, d200p1h3 nanopillars, and 1 kPa hydrogel (n = 20, 20, and 21 cells in flat, nanopillar and hydrogel groups, respectively; mean ± s.e.m). (e) Representative AFM retraction curve for pulling membrane tethers (right) and zoom-in view of a tether rupture event (left) in the retraction curve. The difference in the force before and after the rupture (indicated by green lines) is used to calculate the tether force. (f) Quantitative analysis of tether force obtained from cells on flat quartz, nanopillar quartz, and 1 kPa hydrogel surfaces (n = 21, 21, 22 cells in flat, nanopillar, and hydrogel groups, respectively; mean ± s.e.m.). (g) Fluorescence images of stress fibers (transfected with LifeAct-RFP) in a U2OS cell before and after 20 min of blebbistatin treatment. (h) Young’s modulus of a U2OS cell measured by AFM indentation over the 20 min blebbistatin treatment. *P* value in comparison to flat surfaces was determined by unpaired two-tailed t test (d, f). Scale bars, 10 μm (a, b), 20 μm (g).

We next measured membrane tension by pulling tethers from the cell membrane using a sharp AFM probe. Membrane tethers are lipid nanotubes formed between the AFM probe and the cell surface, which can be stretched up by the AFM probe to produce a constant force until rupture. The tether rupture force is proportional to the square root of the membrane tension^43^ and can be measured as the difference between the force plateaus before and after the rupture (**Figure 2e, Figures S5c–e**). From our measurements, the average tether force for cells on quartz nanopillars was 15% lower than on flat surfaces and is similar to that on 1 kPa hydrogel substrates (**Figure 2f**). These AFM measurements clearly demonstrate that cells on nanopillar quartz and soft hydrogels are much softer than cells on flat quartz surfaces.

Nanotopography has been shown to reduce actin stress fibers.^27, 30^ To confirm the importance of actin fibers in modulating cell mechanics, we continuously monitored the cell stiffness by AFM indentation while the cell was being treated by blebbistatin that disrupts actin stress fibers without affecting cell shape (**Figure 2g**). *In situ* AFM measurement of cells on flat quartz surfaces shows that blebbistatin treatment reduced the cell stiffness in 5 min (**Figure 2h**).

### Nanotopography inhibits YAP activity when cell area and shape are controlled

We found that the presence of nanopillars on quartz substrates drastically decreased cell areas and YAP activity (measured by YAP nucleus/cytosol ratios) (**Figures 3a, b, Figures S6, S7**), agreeing with previous nanotopography studies.^29, 30^ For these measurements, cells were cultured at a low density to reduce cell–cell contact which is known to affect cell size and YAP activity.^44^ Previous studies have demonstrated that reducing cell size was sufficient to inhibit YAP activity.^44–46^ To determine whether nanotopography can affect the YAP activity independently of its effect on cell size, we employed a bioprinting method to precisely control cell size and shape. The bioprinting method uses ultraviolet light-induced cleavage to specify areas that are later coated with cell adhesion molecules such as gelatin.^47^ Unexposed areas are covered with polyethylene glycol to prevent cell adhesion. Hydrogel is not compatible with bioprinting and not included here.

**Figure 3.**
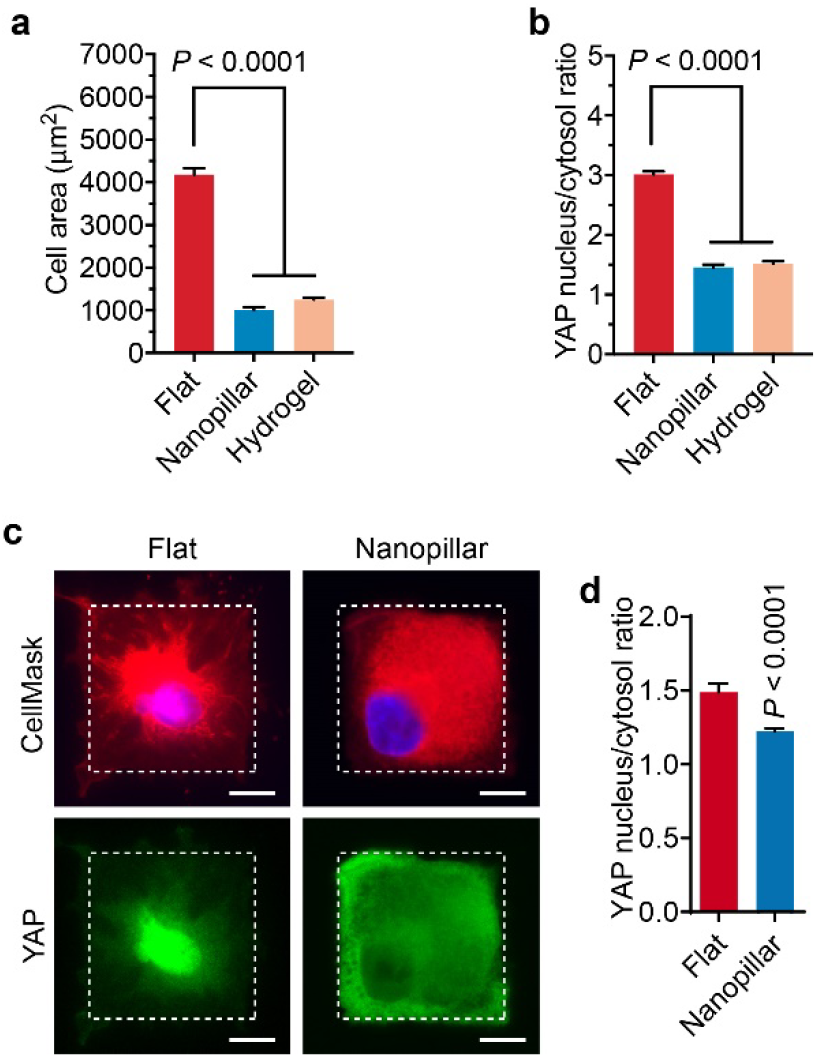
Nanotopgraphy inhibits YAP activity when cell area is controlled. (a) Quantitative analysis of cell area on flat quartz, nanopillar quartz, and 1 kPa hydrogel surfaces. (n= 291, 51, 63 cells in flat, nanopillar, and hydrogel groups, respectively; mean ± s.e.m.). (b) Quantitative analysis of YAP nucleus/cytosol ratio on flat quartz, nanopillar quartz, and 1 kPa hydrogel surfaces. (n= 291, 51, 63 cells in flat, nanopillar, and hydrogel groups, respectively; mean ± s.e.m.). (c) Fluorescence images of hMSCs cultured on flat and d200p1h3 nanopillar surfaces in 2000 μm^2^ square areas confined by bioprinting. CellMask staining (red) reveals the cell shape and YAP localization is indicated with anti-YAP immunostaining (green) and Hoechst staining (blue for nuclei). Dash lines indicate the bioprinting areas. (d) Quantitative analysis of YAP nucleus/cytosol ratio for cells confined to 2000 μm^2^ square areas (n = 14 and 26 cells in flat and nanopillar groups, respectively; mean ± s.e.m.). *P* value in comparison to flat surface was determined by unpaired two-tailed t test. Scale bars: 10 μm (c).

We printed square-shaped areas of 2000 μm^2^ on both flat quartz surfaces and nanopillar quartz surfaces. Co-staining of YAP, the nucleus, and the cell membrane shows that YAP was generally more cytosolic on nanopillars than that on flat quartz surfaces when cell size and shape were the same (**Figures 3c, d**). We note that, when the two-dimensional spreading area was controlled to be the same, cells on nanopillars had a larger total membrane contact area than those on flat surfaces because the plasma membrane wrapped around vertical nanopillars. On flat surfaces, the membrane contact area is positively correlated with YAP activity^41^. The observation that cells on nanopillars had a lower YAP activity despite having larger membrane contact areas further indicates that nanopillars can inhibit YAP activity through mechanisms independent of cell area.

### Nanotopography reduces focal adhesions by enhancing the endocytosis of integrin receptors

To understand the topographic effect at the molecular level, we first confirmed that nanotopography modulates actin stress fibers and focal adhesions as previously reported,^27–29^ by immunostaining key protein components including F-actin, β1, vinculin, paxillin, and phosphorylated focal adhesion kinase (pFAK) (**Figures 4a, b**, **Figure S8**). Actin fibers and focal adhesions are large protein complexes that are primarily responsible for generating and sensing mechanical tension. On flat quartz surfaces, thick and bundled actin fibers were anchored on large focal adhesion patches, identified as elongated patches of integrin β1, vinculin, paxillin, or pFAK. In contrast, on quartz nanopillar surfaces, focal adhesion proteins were either diffusive or appear as small puncta. Actin filaments were much thinner with drastically fewer focal adhesion patches anchored at their ends. The observed reduction is not due to the limited imaging depth when cells are cultured on tall nanopillars, because confocal microscope images projected in z-direction confirmed the same observation (**Figure S9**). We quantified the number of large focal adhesions (>1 μm^2^ in U2OS cells^48^) that are primarily responsible for force generation and substrate stiffness sensing. U2OS cells had about 20 large focal adhesions per cell when cultured on flat quartz surfaces but they lost nearly all large focal adhesions on d200p1h3 quartz (**Figure 4c**, **Figure S8**). This dramatic reduction of focal adhesions and stress fibers on nanopillars likely explains the reduction of cell stiffness on these surfaces.

**Figure 4.**
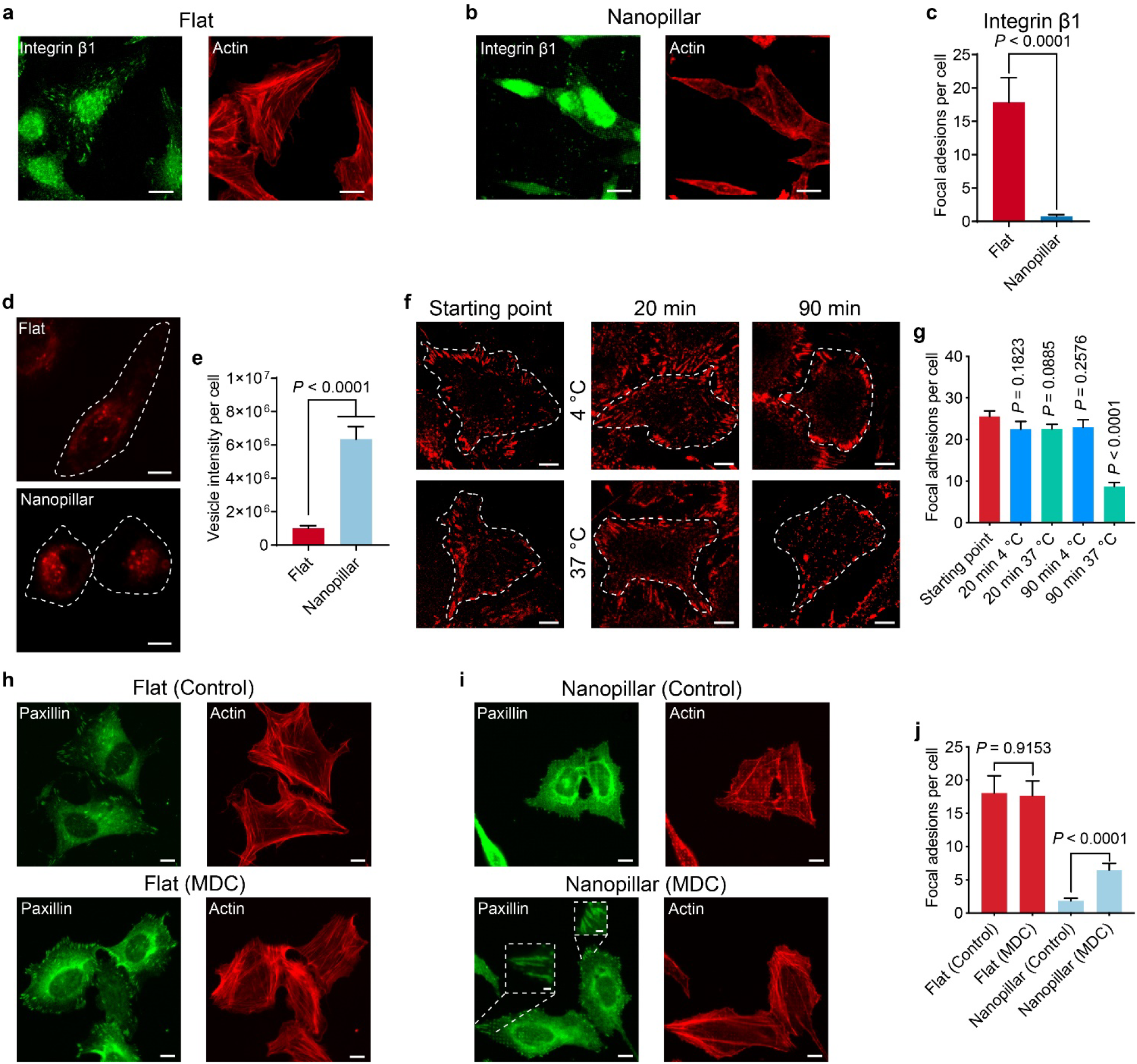
Nanotopography reduces focal adhesions by enhancing the endocytosis of integrin. (a) Fluorescence images of integrin β1 and actin in U2OS cells on nanopillar surfaces. (c) Quantification of large focal adhesion patches (>1 μm^2^) in U2OS cells based on the immunostaining of integrin β1. (d) Fluorescence images of endocytosed FM 1-43 dye in U2OS cells on flat and nanopillar surfaces. The images were obtained by summing the intensity of each pixel of the confocal images over a 12-μm depth with a 500-nm incremental step. (e) Quantitative analysis of endocytic vesicles in U2OS cells on flat quartz and nanopillar quartz surfaces (n = 48 and 30 cells in flat and nanopillar groups, respectively; mean ± s.e.m.). (f) Fluorescence images of integrin β1 affected by temperature-mediated endocytosis. Integrin β1 in U2OS cells was labeled with immunostaining (red). (g) Quantitative analysis of focal adhesion patches (larger than 1 μm^2^) as a result of temperature-mediated endocytosis (n = 10 cells for all groups; mean ± s.e.m.). (h, i) Fluorescence images of paxillin (green) and actin (red) with and without an endocytosis inhibitor MDC treatment, on flat areas (h) and nanopillar areas (i), respectively. (j) Quantitative analysis of focal adhesion patches on flat and nanopillar surfaces before and after MDC treatment (n = 20, 34, 27, and 28 cells in flat (control), flat (MDC), nanopillar (control), nanopillar (MDC) groups, respectively; mean ± s.e.m.). Nanopillars for these experiments had a d200p2.5h1 configuration. *P* values in comparison to flat surface (e), starting point group (g), or control groups (j) were determined by unpaired two-tailed t test. Scale bars, 20 μm (d, f), 4 μm (insets in i).

Integrin receptors play a key role in sensing the substrate stiffness. Clathrin-dependent endocytosis (CME) has been shown to be the primary pathway for removing integrin receptors from cell surface,^49^ which also causes the disassembly of focal adhesions.^50^ For nanotopography, we recently showed that nanotopography enhances clathrin-mediated endocytosis (CME) by locally curving the cell membrane.^51^ Based on these studies, we hypothesize that nanotopography-enhanced endocytosis may be responsible for the reduction of focal adhesions on nanotopography.

We first demonstrate that cells on nanopillar quartz surfaces had enhanced total endocytosis than cells on flat quartz surfaces. To measure endocytosis, cells were incubated with a membrane impermeable FM 1-43 dye for 15 min before the dye was washed out. As it is difficult to discern individual endocytic vesicles in small cells for endocytosis measurements, we used d200p2.5h1 (200 nm diameter, 2.5 μm pitch, and 1 μm height) nanopillars that do not reduce cell size as much as d200p1h3 nanopillars do. Z-stacks of confocal fluorescence images revealed that cells on nanopillar areas had a lot more fluorescence puncta than those on flat quartz surfaces (**Figure 4d**). Quantitative analysis confirms that cells on nanopillars have substantially more endocytosed vesicles labeled with FM 1-43 than those on flat quartz surfaces (**Figure 4e**).

U2OS cells primarily express integrin β1 and β5 isoforms. In this study, substrates were coated with gelatin, which is a binding ligand for integrin β1, but not integrin β5. Therefore, integrin β1 is the primary force receptor for the formation of focal adhesions on the substrates in this study. We followed an established protocol for a pulse-chase experiment to demonstrate that the activation of CME processes causes the removal of integrin β1 from the cell surface and subsequently the disassembly of focal adhesions.^50^ Briefly, cells were maintained at 4 °C to inhibit endocytosis and then stained with an antibody that recognizes the extracellular domain of integrin. Cells were then brought to 37 °C to recover endocytosis for 20 min or 90 min, while control cells were maintained on ice during the same time periods to inhibit endocytosis. We found that cells maintained on ice for 20 or 90 min retained a similar number of focal adhesions as 0 min, whereas cells maintained at 37 °C for 90 min had much fewer large integrin patches with the concurrent appearance of endosome-like spots in the intracellular domain (**Figures 4f, g**). Therefore, the activation of integrin endocytosis results in the disassembly of focal adhesions.

Finally, we demonstrated that blocking CME results in recovery of stress fibers and focal adhesions on nanopillar surfaces. We chose monodansylcadaverine (MDC) as a CME inhibitor. MDC is more specific than other endocytic inhibitors such as PitStop and does not cause observable cell morphology changes.^52^ Cells were treated with either 20 μM MDC or DMSO control for 30 min before they were fixed and stained for actin and paxillin. On flat quartz surfaces, MDC-treated cells and control cells had similar amounts of stress fibers and large focal adhesions (**Figure 4h**). On nanopillars, MDC-treated cells had much more stress fibers and large focal adhesions than the control cells (**Figure 4i**). Quantification of focal adhesions shows that MDC treatment of cells on nanopillars significantly recovered focal adhesions (**Figure 4j**). These results indicate that the enhanced removal of integrin receptors by endocytosis is a key mechanism underlying the reduction of focal adhesions on nanopillar substrates. We noted that MDC treatment only partial recovered focal adhesions, which either indicates that there are additional topography-sensing mechanisms or that our MDC treatment did not sufficiently block integrin endocytosis induced by nanopillars, as MDC concentration is limited by its cytotoxicity.^53^

### Nanotopographical geometry fine-tunes cellular mechanotransduction

To further support the hypothesis that nanotopography modulates mechanotransduction through curvature-mediated endocytosis **(Figure 5a)**, we systematically varied the dimensions of nanopillar arrays. In this study, the nanopillar pitch was varied from 1 to 5 μm, height from 1 to 3 μm, and diameter from 200 to 1000 nm (**Figure 5b, Table S1**). As references to different substrate stiffness, we made a series of polyacrylamide hydrogels ranging from 1 to 14 kPa (**Table S2**).

**Figure 5.**
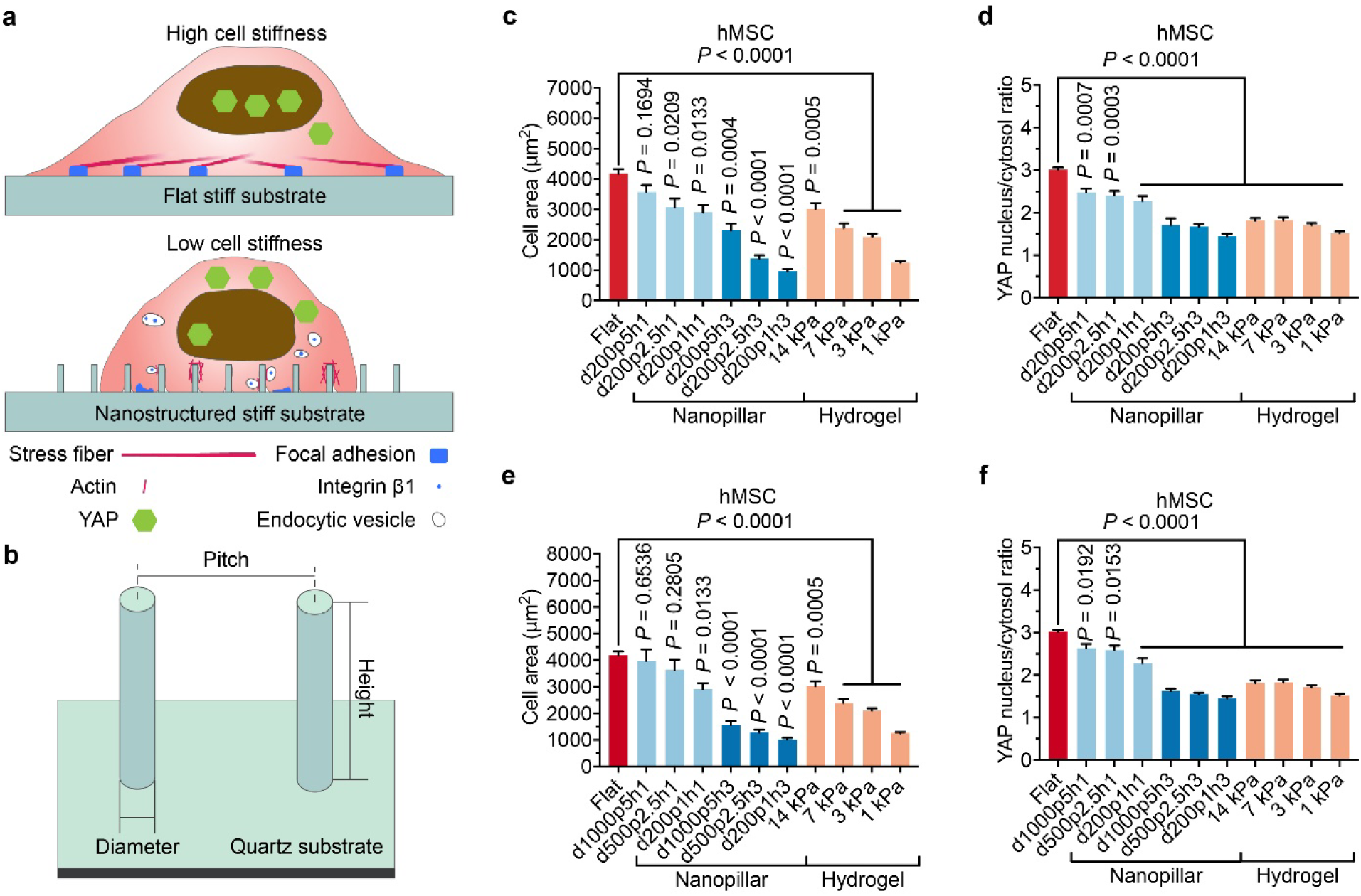
Components of cellular mechanotransduction can be fine-tuned in a wide range by varying nanopillar dimensions. (a) Schematic illustration of focal adhesions and stress fibers in cells cultured on flat and nanopillar areas. (b) Schematic illustration indicating the diameter, pitch, and height of nanopillars. (c, d) Quantitative analyses of cell area (c) and YAP nucleus/cytosol ratio (d) of hMSCs on 200-nm-diameter nanopillars with varied pitches and heights. (n= 15 to 291 cells, Table S3; mean ± s.e.m.). (e, f) Quantitative analyses of cell area (e) and YAP nucleus/cytosol ratio (f) of hMSCs on nanopillars with varied diameters and heights but a constant diameter/pitch ratio. (n= 15 to 291 cells, Table S3; mean ± s.e.m.). *P* value in comparison to flat surface was determined by unpaired two-tailed t test (c-f).

We first varied the curved membrane area by changing the pitch and the height while keeping the nanopillar diameter constant. As shown in **Figure 5c**, the cell areas of hMSCs on flat quartz surfaces were substantially larger than those on hydrogels and quartz nanopillars. For the hydrogel group, cell areas decreased when the hydrogel stiffness was reduced, illustrating the stiffness effect.^6, 38, 41^ For nanopillars of the same height (either 1 or 3 μm), averaged cell areas decreased as the curved membrane area per substrate area was increased by reducing the pitch from 5 to 1 μm (**Figure 5c**). For nanopillars of the same spacing, averaged cell areas decreased as the height of nanopillars increases from 1 to 3 μm to increase curved membrane areas. Furthermore, the YAP nucleus/cytoplasm ratio decreased as the curved membrane area was increased by either reducing the pitch or increasing the height (**Figure 5d**).

Next, we varied nanopillar diameters. We varied the pitch along with the diameter to keep a constant diameter/pitch ratio for efficient membrane wrapping. We found that 200-nm-diameter nanopillars induced a stronger reduction of cell area and YAP activity than 500-nm-diameter and 1000-nm-diamter nanopillars with the same height (**Figures 5e, f**). This observation agrees with our hypothesis based on nanotopography-enhanced CME, since CME is more enhanced by 200-nm-diameter nanopillars than thicker nanopillars^51^. Measurements using U2OS cells revealed similar results as hMSCs (**Figure S10**).

## Conclusions

In this work, we show that nanotopography can drastically modulate cell mechanics such that cells respond to rigid nanopillar substrates (72 GPa) in ways similar to their responses to soft hydrogels (1~14 kPa). We reveal that nanotopography-induced membrane curvatures enhance the endocytosis of integrin receptors, which leads to fewer integrins on the cell surface and subsequently the disassembly of focal adhesions and stress fibers. Previous studies indicate that soft substrates do not provide sufficient resistance to traction forces generated by stress fibers, leading to a lower percentage of activated integrins on the cell surface. Interestingly, although nanotopography initially acts by inducing membrane curvatures and stiffness initially acts through its resistance to the probing force of cells, the two physical cues converge on modulating integrin receptors. These findings will support the rational design of surface topography for such purposes as making cells on a rigid implant behave as if the implant had a stiffness similar to native tissues.

## Associated Content

Nanofabrication and surface functionalization of nanopillar quartz substrates, Preparation and surface functionalization of polyacrylamide hydrogels, cell culture, isolation and culture of hippocampal neurons, staining and imaging of hippocampal neurons, differentiation of stem cells, staining and imaging of stem cells, AFM measurement, staining and imaging of cell area and YAP, bioprinting, staining imaging of focal adhesions and stress fibers, imaging of total endocytosis, temperature-mediated endocytosis inhibition, and drug-induced endocytosis inhibition (PDF)

## Acknowledgements

This work was supported by NIH grants 1R35GM141598 and 1R01GM128142. X.L. thanks the Banting Postdoctoral Fellowships program, administered by the Government of Canada. L.H.K. thanks support from the Carlsberg Foundation. We thank Stanford Nanofabrication Facility and Stanford Nano Shared Facilities for help with nanofabrication; L.Kaplan of B.C. group for teaching dissection; W.Zhao and A.F.McGuire of B.C. group for help with nanofabrication; C.D.Lindsey of S.C.Heilshorn group and A.J.Price of A.R.Dunn group at Stanford for sharing reagents and protocols; Y.Yang of B.C. group and J.Wang of J.Puglisi group for comments on the manuscript; M.Dong at Aarhus University and V.M. Weaver at UC San Francisco for advice on the project. Part of this work was performed at the Stanford Cell Sciences Imaging Facility, supported by Award 1S10OD021514-01 from the National Center for Research Resources.

## Author contributions

B.C., X.L., and L.H.K. conceived the study and designed experiments. X.L., B.C., L.H.K., and T.L.L. wrote the manuscript. X.L. and Z.J. fabricated nanostructures. L.H.K. and C.T. prepared hydrogels. X.L., L.H.K., and W.Z. performed most biological experiments. L.H.K, C.T., and Z.J. performed the bioprinting experiment. X.L. and B.C. designed the software programs for data analysis. All authors discussed the results and commented on the manuscript.

## Competing interests

The authors declare no competing interests.

Correspondence and requests for materials should be addressed to B.C.

## References

1. Engler, A. J.; Sen, S.; Sweeney, H. L.; Discher, D. E., Matrix elasticity directs stem cell lineage specification. Cell 2006, 126 (4), 677–689.

2. Sun, Y. B.; Yong, K. M. A.; Villa-Diaz, L. G.; Zhang, X. L.; Chen, W. Q.; Philson, R.; Weng, S. N.; Xu, H. X.; Krebsbach, P. H.; Fu, J. P., Hippo/YAP-mediated rigidity-dependent motor neuron differentiation of human pluripotent stem cells. Nat Mater 2014, 13 (6), 599–604.

3. Hadden, W. J.; Young, J. L.; Holle, A. W.; McFetridge, M. L.; Kim, D. Y.; Wijesinghe, P.; Taylor-Weiner, H.; Wen, J. H.; Lee, A. R.; Bieback, K.; Vo, B. N.; Sampson, D. D.; Kennedy, B. F.; Spatz, J. P.; Engler, A. J.; Choi, Y. S., Stem cell migration and mechanotransduction on linear stiffness gradient hydrogels. P Natl Acad Sci USA 2017, 114 (22), 5647–5652.

4. Engler, A. J.; Carag-Krieger, C.; Johnson, C. P.; Raab, M.; Tang, H. Y.; Speicher, D. W.; Sanger, J. W.; Sanger, J. M.; Discher, D. E., Embryonic cardiomyocytes beat best on a matrix with heart-like elasticity: scar-like rigidity inhibits beating. J Cell Sci 2008, 121 (22), 3794–3802.

5. Levental, K. R.; Yu, H. M.; Kass, L.; Lakins, J. N.; Egeblad, M.; Erler, J. T.; Fong, S. F. T.; Csiszar, K.; Giaccia, A.; Weninger, W.; Yamauchi, M.; Gasser, D. L.; Weaver, V. M., Matrix crosslinking forces tumor progression by enhancing integrin signaling. Cell 2009, 139 (5), 891–906.

6. Solon, J.; Levental, I.; Sengupta, K.; Georges, P. C.; Janmey, P. A., Fibroblast adaptation and stiffness matching to soft elastic substrates. Biophys J 2007, 93 (12), 4453–4461.

7. Tee, S. Y.; Fu, J. P.; Chen, C. S.; Janmey, P. A., Cell shape and substrate rigidity both regulate cell stiffness. Biophys J 2011, 100 (5), L25–L27.

8. Li, J.; Springer, T. A., Integrin extension enables ultrasensitive regulation by cytoskeletal force. P Natl Acad Sci USA 2017, 114 (18), 4685–4690.

9. Wells, R. G., Tissue mechanics and fibrosis. Bba-Mol Basis Dis 2013, 1832 (7), 884–890.

10. Salatino, J. W.; Ludwig, K. A.; Kozai, T. D. Y.; Purcell, E. K., Glial responses to implanted electrodes in the brain. Nat Biomed Eng 2017, 1 (11), 862–877.

11. Sridharan, A.; Rajan, S. D.; Muthuswamy, J., Long-term changes in the material properties of brain tissue at the implant-tissue interface. J Neural Eng 2013, 10 (6).

12. Lacour, S. P.; Courtine, G.; Guck, J., Materials and technologies for soft implantable neuroprostheses. Nat Rev Mater 2016, 1 (10).

13. Jacot, J. G.; McCulloch, A. D.; Omens, J. H., Substrate stiffness affects the functional maturation of neonatal rat ventricular myocytes. Biophys J 2008, 95 (7), 3479–3487.

14. Tanaka, A.; Fujii, Y.; Kasai, N.; Okajima, T.; Nakashima, H., Regulation of neuritogenesis in hippocampal neurons using stiffness of extracellular microenvironment. Plos One 2018, 13 (2).

15. Zhang, Y. S.; Khademhosseini, A., Advances in engineering hydrogels. Science 2017, 356 (6337).

16. Gao, X.; Fraulob, M.; Haiat, G., Biomechanical behaviours of the bone-implant interface: a review. J R Soc Interface 2019, 16 (156).

17. Smeets, R.; Stadlinger, B.; Schwarz, F.; Beck-Broichsitter, B.; Jung, O.; Precht, C.; Kloss, F.; Grobe, A.; Heiland, M.; Ebker, T., Impact of dental implant surface modifications on osseointegration. Biomed Res Int 2016, 2016.

18. Merrill, D. R., Materials considerations of implantable neuroengineering devices for clinical use. Curr Opin Solid St M 2014, 18 (6), 329–336.

19. Chen, W. Q.; Villa-Diaz, L. G.; Sun, Y. B.; Weng, S. N.; Kim, J. K.; Lam, R. H. W.; Han, L.; Fan, R.; Krebsbach, P. H.; Fu, J. P., Nanotopography influences adhesion, spreading, and self-renewal of human embryonic stem cells. Acs Nano 2012, 6 (5), 4094–4103.

20. Ding, Y. F.; Sun, J. R.; Ro, H. W.; Wang, Z.; Zhou, J.; Lin, N. J.; Cicerone, M. T.; Soles, C. L.; Lin-Gibson, S., Thermodynamic underpinnings of cell alignment on controlled topographies. Adv Mater 2011, 23 (3), 421–425.

21. Jacchetti, E.; Di Rienzo, C.; Meucci, S.; Nocchi, F.; Beltram, F.; Cecchini, M., Wharton’s Jelly human Mesenchymal Stem Cell contact guidance by noisy nanotopographies. Sci Rep-Uk 2014, 4.

22. Sun, J. R.; Ding, Y. F.; Lin, N. J.; Zhou, J.; Ro, H.; Soles, C. L.; Cicerone, M. T.; Lin-Gibson, S., Exploring cellular contact guidance using gradient nanogratings. Biomacromolecules 2010, 11 (11), 3067–3072.

23. Kwon, K. W.; Park, H.; Song, K. H.; Choi, J. C.; Ahn, H.; Park, M. J.; Suh, K. Y.; Doh, J., Nanotopography-guided migration of T cells. J Immunol 2012, 189 (5), 2266–2273.

24. Wu, Y. N.; Law, J. B. K.; He, A. Y.; Low, H. Y.; Hui, J. H. P.; Lim, C. T.; Yang, Z.; Lee, E. H., Substrate topography determines the fate of chondrogenesis from human mesenchymal stem cells resulting in specific cartilage phenotype formation. Nanomed-Nanotechnol 2014, 10 (7), 1507–1516.

25. Teo, B. K. K.; Wong, S. T.; Lim, C. K.; Kung, T. Y. S.; Yap, C. H.; Ramagopal, Y.; Romer, L. H.; Yim, E. K. F., Nanotopography modulates mechanotransduction of stem cells and induces differentiation through focal adhesion kinase. Acs Nano 2013, 7 (6), 4785–4798.

26. Antonini, S.; Meucci, S.; Parchi, P.; Pacini, S.; Montali, M.; Poggetti, A.; Lisanti, M.; Cecchini, M., Human mesenchymal stromal cell-enhanced osteogenic differentiation by contact interaction with polyethylene terephthalate nanogratings. Biomed Mater 2016, 11 (4).

27. Lou, H. Y.; Zhao, W. T.; Li, X.; Duan, L. T.; Powers, A.; Akamatsu, M.; Santoro, F.; McGuire, A. F.; Cui, Y.; Drubin, D. G.; Cui, B. X., Membrane curvature underlies actin reorganization in response to nanoscale surface topography. P Natl Acad Sci USA 2019, 116 (46), 23143–23151.

28. Kuo, C. W.; Chueh, D. Y.; Chen, P. L., Investigation of size-dependent cell adhesion on nanostructured interfaces. J Nanobiotechnol 2014, 12.

29. Loye, A. M.; Kinser, E. R.; Bensouda, S.; Shayan, M.; Davis, R.; Wang, R.; Chen, Z.; Schwarz, U. D.; Schroers, J.; Kyriakides, T. R., Regulation of mesenchymal stem cell differentiation by nanopatterning of bulk metallic glass. Sci Rep-Uk 2018, 8.

30. Hansel, C. S.; Crowder, S. W.; Cooper, S.; Gopal, S.; da Cruz, M. J. P.; Martins, L. D.; Keller, D.; Rothery, S.; Becce, M.; Cass, A. E. G.; Bakal, C.; Chiappini, C.; Stevens, M. M., Nanoneedle-mediated stimulation of cell mechanotransduction machinery. Acs Nano 2019, 13 (3), 2913–2926.

31. Song, L. Q.; Wang, K.; Li, Y.; Yang, Y., Nanotopography promoted neuronal differentiation of human induced pluripotent stern cells. Colloid Surface B 2016, 148, 49–58.

32. Li, X.; Matino, L.; Zhang, W.; Klausen, L.; McGuire, A. F.; Lubrano, C.; Zhao, W. T.; Santoro, F.; Cui, B. X., A nanostructure platform for live-cell manipulation of membrane curvature. Nat Protoc 2019, 14 (6), 1772–1802.

33. Fu, J. P.; Wang, Y. K.; Yang, M. T.; Desai, R. A.; Yu, X. A.; Liu, Z. J.; Chen, C. S., Mechanical regulation of cell function with geometrically modulated elastomeric substrates. Nat Methods 2010, 7 (9), 733–U95.

34. Santoro, F.; Zhao, W. T.; Joubert, L. M.; Duan, L. T.; Schnitker, J.; van de Burgt, Y.; Lou, H. Y.; Liu, B. F.; Salleo, A.; Cui, L. F.; Cui, Y.; Cui, B. X., Revealing the cell-material interface with nanometer resolution by focused ion beam/scanning electron microscopy. Acs Nano 2017, 11 (8), 8320–8328.

35. Barnes, A. P.; Polleux, F., Establishment of axon-dendrite polarity in developing neurons. Annu Rev Neurosci 2009, 32, 347–381.

36. Chang, T. Y.; Chen, C.; Lee, M.; Chang, Y. C.; Lu, C. H.; Lu, S. T.; Wang, D. Y.; Wang, A.; Guo, C. L.; Cheng, P. L., Paxillin facilitates timely neurite initiation on soft-substrate environments by interacting with the endocytic machinery. Elife 2017, 6.

37. Wen, J. H.; Vincent, L. G.; Fuhrmann, A.; Choi, Y. S.; Hribar, K. C.; Taylor-Weiner, H.; Chen, S. C.; Engler, A. J., Interplay of matrix stiffness and protein tethering in stem cell differentiation. Nat Mater 2014, 13 (10), 979–987.

38. Qian, W. Y.; Gong, L. Q.; Cui, X.; Zhang, Z. J.; Bajpai, A.; Liu, C.; Castillo, A. B.; Teo, J. C. M.; Chen, W. Q., Nanotopographic regulation of human mesenchymal stem cell osteogenesis. Acs Appl Mater Inter 2017, 9 (48), 41794–41806.

39. Abagnale, G.; Steger, M.; Nguyen, V. H.; Hersch, N.; Sechi, A.; Joussen, S.; Denecke, B.; Merkel, R.; Hoffmann, B.; Dreser, A.; Schnakenberg, U.; Gillner, A.; Wagner, W., Surface topography enhances differentiation of mesenchymal stem cells towards osteogenic and adipogenic lineages. Biomaterials 2015, 61, 316–326.

40. Ahn, E. H.; Kim, Y.; Kshitiz; An, S. S.; Afzal, J.; Lee, S.; Kwak, M.; Suh, K. Y.; Kim, D. H.; Levchenko, A., Spatial control of adult stem cell fate using nanotopographic cues. Biomaterials 2014, 35 (8), 2401–2410.

41. Yeung, T.; Georges, P. C.; Flanagan, L. A.; Marg, B.; Ortiz, M.; Funaki, M.; Zahir, N.; Ming, W. Y.; Weaver, V.; Janmey, P. A., Effects of substrate stiffness on cell morphology, cytoskeletal structure, and adhesion. Cell Motil Cytoskel 2005, 60 (1), 24–34.

42. Vargas-Pinto, R.; Gong, H.; Vahabikashi, A.; Johnson, M., The effect of the endothelial cell cortex on atomic force microscopy measurements. Biophys J 2013, 105 (2), 300–309.

43. Krieg, M.; Helenius, J.; Heisenberg, C. P.; Muller, D. J., A bond for a lifetime: Employing membrane nanotubes from living cells to determine receptor-ligand kinetics. Angew Chem Int Edit 2008, 47 (50), 9775–9777.

44. Dupont, S.; Morsut, L.; Aragona, M.; Enzo, E.; Giulitti, S.; Cordenonsi, M.; Zanconato, F.; Le Digabel, J.; Forcato, M.; Bicciato, S.; Elvassore, N.; Piccolo, S., Role of YAP/TAZ in mechanotransduction. Nature 2011, 474 (7350), 179–83.

45. von Erlach, T. C.; Bertazzo, S.; Wozniak, M. A.; Horejs, C. M.; Maynard, S. A.; Attwood, S.; Robinson, B. K.; Autefage, H.; Kallepitis, C.; Hernandez, A. D.; Chen, C. S.; Goldoni, S.; Stevens, M. M., Cell-geometry-dependent changes in plasma membrane order direct stem cell signalling and fate. Nat Mater 2018, 17 (3), 237-+.

46. Kilian, K. A.; Bugarija, B.; Lahn, B. T.; Mrksich, M., Geometric cues for directing the differentiation of mesenchymal stem cells. P Natl Acad Sci USA 2010, 107 (11), 4872–4877.

47. Strale, P. O.; Azioune, A.; Bugnicourt, G.; Lecomte, Y.; Chahid, M.; Studer, V., Multiprotein printing by light-induced molecular adsorption. Adv Mater 2016, 28 (10), 2024-+.

48. Legerstee, K.; Geverts, B.; Slotman, J. A.; Houtsmuller, A. B., Dynamics and distribution of paxillin, vinculin, zyxin and VASP depend on focal adhesion location and orientation. Sci Rep-Uk 2019, 9.

49. Bridgewater, R. E.; Norman, J. C.; Caswell, P. T., Integrin trafficking at a glance. J Cell Sci 2012, 125 (16), 3695–3701.

50. Chao, W. T.; Kunz, J., Focal adhesion disassembly requires clathrin-dependent endocytosis of integrins. Febs Lett 2009, 583 (8), 1337–1343.

51. Zhao, W. T.; Hanson, L.; Lou, H. Y.; Akamatsu, M.; Chowdary, P. D.; Santoro, F.; Marks, J. R.; Grassart, A.; Drubin, D. G.; Cui, Y.; Cui, B. X., Nanoscale manipulation of membrane curvature for probing endocytosis in live cells. Nat Nanotechnol 2017, 12 (8), 750-+.

52. Guo, S.; Zhang, X.; Zheng, M.; Zhang, X.; Min, C.; Wang, Z.; Cheon, S. H.; Oak, M. H.; Nah, S. Y.; Kim, K. M., Selectivity of commonly used inhibitors of clathrin-mediated and caveolae-dependent endocytosis of G protein-coupled receptors. Bba-Biomembranes 2015, 1848 (10), 2101–2110.

53. Gilad, G. M.; Gilad, V. H., Cytotoxic effects of monodansylcadaverine and methylamine in primary cultures of rat cerebellar neurons. Int J Dev Neurosci 1986, 4 (5), 401–5.

