## Supporting Information for "Nanoscale surface topography reduces focal adhesions and cell stiffness by enhancing integrin endocytosis"

### Methods

**Nanofabrication and surface functionalization of nanopillar quartz substrates.** Nanostructures on quartz substrates were fabricated with a protocol similar to our previous publication.^1^ Briefly, a quartz wafer (Silicon Materials, cat. no. 04Q 525-25-1F-SO) was cut into pieces and coated with a layer of electron beam resist (AllResist, cat. no. AR-P 6200) by spin coating. An E-beam lithography (JEOL, model no. JBX-6300FS) system was used to expose desired patterns, and then the pieces were developed. A 120 nm thick chromium layer was deposited to the chips and then lifted off to keep only the Cr patterns. C_4_F_8_-based dry etching was used to make nanopillars with the desired height. At the end of nanofabrication, Cr masks on top of nanostructures were removed by an etchant, and the chips were cleaned in a heated piranha solution. The nanotopographical configurations used in this study are listed in **Table S1**.

Quartz nanopillar chips were functionalized with a covalent bonding-based surface chemistry process for stable attachment of U2OS cells and human mesenchymal stem cells (hMSCs). After each chip (surface area 1 cm^2^) was treated by a plasma cleaner (Harrick, cat. no. PDC-3XG), a 10-µl drop of (3-aminopropyl)triethoxysilane (APTES; Sigma-Aldrich, cat. no. A3648) was applied to the chip surface for 5 min, followed by extensive rinsing in deionized water and 30 min of incubation in 0.5% glutaraldehyde (Sigma-Aldrich, cat. no. P5882). The chip was rinsed again in deionized water and then incubated upside-down on a drop of 0.1% (wt/vol) gelatin (Sigma-Aldrich, cat. no. G9391) on parafilm for 1 hr at room temperature. The chip was washed three times in phosphate-buffered saline (PBS) and kept in PBS until cell culture. Coated chips were sterilized with ultraviolet light before cell culture. For hippocampal neuron culture, the chip was cleaned with air plasma and then its surface was incubated with 0.1 mg ml^–1^ poly-L-lysine (PLL, Sigma-Aldrich, cat. No. P6282) and 50 µg ml^–1^ rat tail collagen I (Corning, cat. no. 354236).

**Preparation and surface functionalization of polyacrylamide hydrogels.** A 1 cm diameter glass coverslip (Fisher Scientific, cat. no. CB00100RA020MNT0) was cleaned in ethanol, followed by air plasma cleaning for 30 min (Harrick, cat. no. PDC-3XG). A 10-ul drop of APTES was applied to the coverslip surface for 5 min. This was followed by extensive rinsing in deionized water and 30 min of incubation in 0.5% (vol/vol) glutaraldehyde. A 989-µl gel solution was obtained by mixing acrylamide monomer (Fisher Scientific, cat. no. AC164851000), *N,N* methylene-bis-acrylamide crosslinker (Fisher Scientific, cat. no. AC163800050) and milli-Q water at the indicated concentrations (**Table S2**). The solution was placed in a desiccator and degassed for 5 min to remove oxygen from the solution. The polymerization reaction was then initiated by adding 10 µl 10% (wt/vol) ammonium persulfate (APS; Sigma-Aldrich, cat. no. 248614) and 1 µl *N,N,N’,N’*-tetramethylethylenediamine (TEMED; Fisher Scientific, cat. no. 89-202), and the solution was carefully mixed by pipetting without introducing air bubbles. A 12-µl solution was then immediately added to a clean coverslip placed in a petri dish, and the functionalized coverslip was placed upside-down on top of the drop. The remaining gel solution was used to observe the polymerization progress. The polymerization process usually takes less than 10 min. When the gel solution was solidified, the petri dish was filled with PBS and incubated for an additional 10 min. The coverslips were then carefully separated using a scalpel knife and the coverslip with hydrogel stored in PBS.

To treat the hydrogel surface for cell attachment, the hydrogel was incubated with 1 mg ml^–1^ Sulfo-SANPAH (Sigma-Aldrich, cat. no. 803332), activated with ultraviolet light for 7 min, and washed in PBS. The incubation, activation, and washing steps were repeated a second time. The hydrogel was then incubated upside-down on a drop of 0.1% (wt/vol) gelatin (Sigma-Aldrich, cat. no. G9391) on parafilm for 1 hr at room temperature. Coated hydrogels were sterilized with ultraviolet light before cell culture. For hippocampal neuron culture, the hydrogel samples were incubated with 0.1 mg ml^–1^ of poly-L-lysine (PLL) and 50 µg ml^–1^ collagen. Mechanical characterization of hydrogels was done using an atomic force microscope (AFM; Bruker, model no. BioScope Resolve). Cantilevers with nominal spring constants of 0.35 N m^–1^ were calibrated using the thermal noise method and 20 µm polystyrene beads (Microparticles GmbH, cat. no. PS-R-20.0) were glued to the tip of the cantilever using the UV curable glue Norland Optical Adhesive 63 (Fisher Scientific, cat. no. NC9936552). The hydrogels were then characterized in PBS at 37 °C using 16 indentation curves with force controlled to reach approximately 1 µm indentation. The data was analyzed in Nanoscope Analysis 1.9 (Bruker) by baseline correcting each approach curve and fitting the data to the Hertz model using the best estimate fit for the contact point and a Poisson’s ratio of 0.5.^50^

**Cell culture.** U2OS cells (ATCC, cat. no. HTB-96) were cultured in Dulbecco’s Modified Eagle Medium (DMEM; Sigma-Aldrich, cat. no. D6546) supplemented with 10% (vol/vol) fetal bovine serum (FBS; Sigma-Aldrich, cat. no. F7524) and 1% (vol/vol) penicillin-streptomycin (Sigma-Aldrich, cat. no. P4333). Embryonic hippocampal neurons were cultured in Neurobasal Medium (Fisher Scientific, cat. no. 21103-49) supplemented with 2% (vol/vol) B27 (Gibco, cat. no. 17504044), 1% (vol/vol) glutamax (Gibco, cat. no. 35050061), and 100 unit ml^–1^ penicillin-streptomycin. Human mesenchymal stem cells (hMSCs; bone marrow-derived, RoosterBio, cat. no. MSC-003) were cultured in the culture medium included in the kit (RoosterBio, cat. no. KT-002).

**Isolation and culture of hippocampal neurons.** Primary hippocampal neuron cells were dissected from E18 rat embryos as previously described.^2^ The isolated hippocampi were pooled and dissociated by incubation in 5 ml of 0.25% trypsin for 30 minutes at 37 °C followed by quenching and mechanical dissociation by trituration. The cell suspension was spun down and resuspended to a density of 3 × 10^5^ cells ml^–1^ in Neurobasal medium with 2% (vol/vol) B27. 100 μl of the suspension was added to each nanopillar chip or hydrogel, which was then incubated for 2 hr to allow cell attachment before transferring to a 24 well plate with 1 ml Neurobasal medium with 2% (vol/vol) B27. The cells were incubated for a total of 20 hr before further processing. Rats used in this study were treated in accordance with Stanford University’s institutional guidelines.

**Staining and imaging of hippocampal neurons.** For quantifying neurite outgrowth, hippocampal neurons were first fixed with 4% paraformaldehyde for 10 min followed by permeabilized in 0.1% Triton X-100 for 10 min and blocking in 1% BSA for 30 minutes. The cells were then stained with anti-tau-1 antibody produced in mouse (EMD Millipore, cat. no. MAB3420) and anti-MAP2 antibody produced in rabbit (Sigma-Aldrich, cat. no. M3696) for 30 min. The cells were finally stained with anti-rabbit antibody coupled to Alexa 488 (Invitrogen, cat. no. A27034), anti-mouse antibody coupled to Alexa 594 (Invitrogen, cat. no. R37121) and Hoechst (Sigma-Aldrich, cat. no. 94403) for 30 min. All staining steps were performed at room temperature. The cells were imaged on an epifluorescence microscope (Leica, model no. DMI6000 B) using a 20× objective, and images were analyzed in ImageJ using a custom script and the Simple Neurite Tracer plugin. Neurite lengths were measured from the anti-MAP2 staining by tracing each neurite from the end to the main cell body. Cells with at least one neurite longer than 35 μm were counted. 207, 193, and 177 cells were analyzed for flat quartz, d200p1h3 nanopillars, and 1 kPa hydrogel, respectively.

**Differentiation of stem cells.** hMSCs with a passage number from three to five were used in differentiation. Gibco™ StemPro™ Osteogenesis Differentiation Kit (Gibco, cat. no. A1007201) and Adipogenesis Differentiation Kit (Gibco, cat. no. A1007001) were purchased from Fisher Scientific and prepared as instructed. For osteogenic differentiation, hMSCs were plated at a density of 2 × 10^3^ cm^–2^ and cultured for 14 days with the osteogenic differentiation kit. For osteogenic differentiation, hMSCs were plated at a density of 4 × 10^3^ cm^–2^ and cultured for 14 days with the osteogenic differentiation kit. For adipogenic differentiation, hMSCs were plated at a density of 2.5 × 10^4^ cm^–2^ in regular growth medium and grown to reach confluency, before the medium was replaced with the adipogenic differentiation kit and grown for another six days.

**Staining and imaging of stem cells.** After differentiation, hMSCs were fixed in 4% (wt/vol) paraformaldehyde and washed with PBS, followed by staining with an osteogenic or adipogenic marker. For osteogenic staining, leukocyte alkaline phosphatase (ALP) kit (Sigma-Aldrich, cat. no. 86C) was applied according to the instructions. For adipogenic staining, Oil Red O (Sigma-Aldrich, cat. no. O0625) was used to label the lipids. Oil Red O was dissolved in isopropanol to reach a concentration of 0.5% (wt/vol) as a stock solution, which was later dissolved in MilliQ water in a 3:2 ratio to obtain a stain solution. The stain solution was made fresh and filtered. The samples were soaked with the stain solution for 20 min, before thoroughly washed with MilliQ water. All the samples of both osteogenic and adipogenic differentiation were stained with Hoechst to label nuclei. The stained samples were imaged with a true-color microscope (Keyence, model no. BZX-800). The obtained images were prepared with ImageJ and analyzed with an automated Matlab code.

In the analysis of osteogenic differentiation, the number of cells in each image was calculated by Hoechst staining, and the positive ones were counted by the ALP staining intensity in the nuclei. The number of stained cells was divided by the total number of cells in each image, to obtain an osteogenic differentiation rate. In the analysis of adipogenic differentiation, the number of cells in each image was also calculated by Hoechst staining. The average area of cells was obtained by measuring random cells stained by Oil Red O. In each image, the total Oil Red O stained area was measured and divided by the total cell area (average area of cells × total cell amount in the image), to obtain an adipogenic differentiation rate.

**Atomic force microscopy measurement.** U2OS cells were seeded on quartz chips or hydrogels at a density of 1 × 10^4^ cells cm^–2^ and cultured overnight (16 to 20 hr). The chips and gels were then moved to a petri dish with pre-equilibrated medium and loaded on an atomic force microscope (AFM; Bruker, model no. BioScope Resolve). The AFM was operated with a heating stage set to 37 °C and a perfusion cell providing 5% CO_2_. The setup was optimized for live-cell imaging, and no change in cell morphology was observed during the measurements. All AFM characterization of cells was carried out using soft silicon nitride cantilevers (Bruker, cat. no. MLCT-BIO-DC). The cell topography was first imaged using sharp AFM probes with a nominal tip radius of 20 nm and a nominal spring constant of 0.01 N m^–1^. 75 µm^2^ images of 256 × 256 pixels were captured with a scan rate of 0.25 Hz and an applied force of approximately 0.3 nN. This force was high enough to reveal cytoskeletal structures under the membrane but not so high that the membrane would be damaged. The apparent cell stiffness was measured using bead-modified AFM probes. 20 μm diameter polystyrene beads (Microparticles GmbH, cat. no. PS-R-20.0) were glued to the tip of cantilevers with a nominal spring constant of 0.07 N m^–1^ using the UV curable glue Norland Optical Adhesive 63 (Fisher Scientific, cat. No. NC9936552). The cantilever spring constants were calibrated before bead attachment by a thermal noise method. The force-volume mode was used to measure the Young’s modulus of cells. In short, we aligned the AFM probe to the center of the cell using a 20× objective and then completed a 50 μm^2^ force volume image of 20 × 20 pixels. The approach and retract velocities were controlled at 10 μm s^–1^, and the force threshold set to 2 nN. The data was analyzed in Nanoscope Analysis 1.9 (Bruker) by selecting points on the flat top part of the cell, baseline correcting each approach curve and fitting the data within 10~90% force region to the Hertz model using a best estimate fit for the contact point and a Poisson’s ratio of 0.5.^3^

Membrane tether pulling was done in a similar fashion using an AFM probe with a nominal tip radius of 20 nm and a nominal spring constant of 0.01 N m^–1^. The probe was air plasma treated before use to improve tether formation. The tip was aligned to the center of a cell and a 10 μm^2^ force volume image of 4 × 4 pixels was produced. Each force-displacement curve consisted of a 2 μm s^–1^ approach step until a force of 1 nN was reached, then a 0.5 s hold time, and finally a 1 μm s^–1^ retract step of at least 5 μm. The data were analyzed using a custom-written Matlab program (Mathworks) with the NanoScope file I/O utilities package. In short, the derivative of the retraction curve was used to identify positions with a sudden jump in force (tether rupture), and the tether force was calculated by fitting line profiles to the region immediately before and after the force jump. Events within the contact region of the retract curve were disregarded to avoid counting adhesion events as tethers. Force-displacement curves with multiple tethers were also included in the data analysis, but tether events occurring within 100 nm from each other were disregarded (see Figure S5 for an example of a retract curve with multiple membrane tethers). On average, 16, 15, and 24 tether events were recorded for each cell on flat quartz, d200p1h3 nanopillars and 1 kPa hydrogel, respectively.

**Staining and imaging of cell area and YAP.** After culture on a substrate, cells were fixed (4% (wt/vol) paraformaldehyde in PBS), permeabilized (0.1% (vol/vol) Triton X in PBS) and blocked (1% (vol/vol) Triton X and 1% (wt/vol) BSA in PBS). Subsequently, CellMask Orange (ThermoFisher Scientific, cat. no. C10045) was diluted to 5 μg ml^–1^, and a 50 μl of the working solution was applied to each sample to stain the plasma membrane for the measurement of cell area. For YAP staining, anti-YAP antibody produced in mouse (63.7) (Santa Cruz Biotechnology, cat. no. sc-101199) was applied to each chip and incubated for 30 min. After washing, anti-mouse secondary antibody coupled with Alexa 594 (Invitrogen, cat. no. R37121) and Hoechst were then applied to each chip. Samples were imaged with a fully automated inverted epifluorescence microscope (Leica, model no. DMI6000 B). Obtained images were analyzed in a custom-written Matlab program. CellMask and Hoechst images were analyzed in the program to determine cell area and nuclear area, respectively. Cell area and nuclear area were used as masks for calculating YAP nucleus/cytosol ratios.

**Bioprinting.** Quartz nanopillar chips were treated with a plasma cleaner and incubated with PLL grafted with polyethylene glycol (PLL(20)-g[3.5]-PEG(2)) (SuSoS AG, Dübendorf) for 1 hr at room temperature. The solution was aspirated, and the chips were washed 10 times with deionized water without drying the surface before adding the photo-initiator PLPP (Alvéole). Micropatterning was then performed using an optical module (Alvéole, model no. PRIMO) with 2000 mJ mm^–2^ exposure, followed by ten rinses with deionized water and 5 min of incubation with 0.1% (vol/vol) gelatin-FITC (BioVision, cat. no. M1303) at room temperature. The chips were finally washed in PBS before trypsinized cells were added to a final density of 5 × 10^4^ cells cm^–2^ and the chips incubated for 1 hr. The media was replaced, and the chips incubated overnight before fixation and staining.

**Staining and imaging of focal adhesions and stress fibers.** After culture on a substrate, cells were fixed (4% (wt/vol) paraformaldehyde in PBS), permeabilized (0.1% (vol/vol) Triton X in PBS) and blocked (1% (vol/vol) Triton X and 1% (wt/vol) BSA in PBS). They were then stained with anti-integrin β1 antibody (Sigma-Aldrich, cat. no. MAB2079Z), anti-vinculin antibody (Sigma-Aldrich, cat. no. V9131), anti-paxillin antibody (BD Transduction Laboratory, 610051), or anti-pFAK (phospho Y397, Abcam, cat. no. ab81298), followed by staining of a corresponding secondary antibody with Alexa 488. Cells were also stained with Alexa Fluor 647 Phalloidin (Fisher Scientific, cat. no. A22287) and Hoechst. Samples were imaged with a confocal microscope (Nikon, model no. A1R). Vinculin images were analyzed in a custom-written Matlab program which counted the number of focal adhesion patches (larger than 1 μm^2^ and 1.5 μm^2^ for U2OS cells and hMSCs, respectively) in each cell.

**Imaging of total endocytosis.** U2OS cells were cultured on a substrate for 24 hr. The culture medium was removed, and the cells were washed with PBS. Then a 1 ml volume of 2 μM FM 1-43 dye (fixable; ThermoFisher, cat. no. T35356) was added to each 24-well, and the cells were incubated for 15 min in an incubator (Thermo Fisher Scientific, model no. Forma). The dye solution was then removed, and cells were incubated with 1ml PBS for 20 min. The samples were then fixed and stained with Hoechst. The samples were imaged with a confocal microscope, with an incremental 500 nm step in a ±6 um z-range. The obtained images were analyzed in a custom-written Matlab program, where FM 1-43 dye puncta in each focal plane were measured, and the total intensity of the puncta was calculated in each cell.

**Temperature-mediated endocytosis inhibition.** The effect of endocytosis on β1 integrin internalization was measured using an established protocol.^4^ U2OS cells were seeded at a density of 1 × 10^4^ cells cm^–2^ on gelatin-coated coverslips and incubated for 20 hr before serum starving overnight. The cells were then incubated with 10 μM nocodazole in serum-free medium for 1 hr at 37 °C, washed in cold PBS and incubated with anti-integrin β1 antibody for 30 minutes on ice, washed three times with cold PBS and incubated with anti-mouse antibody coupled to Alexa 594 for 30 min on ice. The cells were washed 3 times with cold PBS and incubated in complete media at 37 °C or on ice. The incubation time was 0 (control), 20 or 90 min, after which the cells were washed 3 times with cold PBS. The cells were immediately imaged by fluorescence microscopy, and large integrin patches or focal adhesions counted.

**Drug-induced endocytosis inhibition.** 20 μM MDC (Sigma, cat. no. 30432) was prepared in serum-free DMEM with 0.5% (wt/wt) dimethyl sulfoxide (DMSO). After cells were cultured on a substrate for 24 hr, the culture medium was replaced with this MDC medium. Cells were incubated for 30 min in an incubator, and then washed three times with PBS. Cells in the control group were treated the same with serum-free DMEM containing only 0.5% DMSO.

**Statistical analysis.** All tests were performed using Prism (GraphPad Software) and data are presented as mean ± s.e.m. Statistical significance was evaluated with unpaired two-tailed t test. All experiments were repeated at least three times, unless otherwise stated in the figure captions.

**References**

1.	Li, X.; Matino, L.; Zhang, W.; Klausen, L.; McGuire, A. F.; Lubrano, C.; Zhao, W. T.; Santoro, F.; Cui, B. X., A nanostructure platform for live-cell manipulation of membrane curvature. *Nat Protoc* **2019,** *14* (6), 1772-1802.

2.	Dotti, C. G.; Sullivan, C. A.; Banker, G. A., The establishment of polarity by hippocampal neurons in culture. *J Neurosci* **1988,** *8* (4), 1454-68.

3.	Lin, D. C.; Dimitriadis, E. K.; Horkay, F., Robust strategies for automated AFM force curve analysis - I. Non-adhesive indentation of soft, inhomogeneous materials. *J Biomech Eng-T Asme* **2007,** *129* (3), 430-440.

4.	Chao, W. T.; Kunz, J., Focal adhesion disassembly requires clathrin-dependent endocytosis of integrins. *Febs Lett* **2009,** *583* (8), 1337-1343.

**Table S1.** Configurations of nanostructures.

| **Configuration name** | **Diameter (nm)** | **Pitch (μm)** | **Height (μm)** |
| --- | --- | --- | --- |
| **d1000p2.5h1** | 1000 | 2.5 | 1 |
| **d500p2.5h1** | 500 | 2.5 | 1 |
| **d200p1h1** | 200 | 1 | 1 |
| **d200p2.5h1** | 200 | 2.5 | 1 |
| **d200p5h1** | 200 | 5 | 1 |
| **d200p1h3** | 200 | 1 | 3 |
| **d200p2.5h3** | 200 | 2.5 | 3 |
| **d200p5h3** | 200 | 5 | 3 |

**Table S2.** Configurations of polyacrylamide hydrogels.

| **Configuration name** | **Gel concentration (%; vol/vol)** | **40% acrylamide monomer (μl)** | **2% acrylamide crosslinker (μl)** | **Milli-Q water (μl)** | **TEMED (μl)** | **10% APS (μl)** | **Young’s modulus (kPa; n =16 for all groups; mean ± s.e.m.)** |
| --- | --- | --- | --- | --- | --- | --- | --- |
| 1 kPa | 4 | 100 | 40 | 849 | 1 | 10 | 1.039 ± 0.0105 |
| 3 kPa | 5 | 125 | 50 | 814 | 1 | 10 | 2.824 ± 0.0179 |
| 7 kPa | 7 | 175 | 70 | 744 | 1 | 10 | 6.871 ± 0.0395 |
| 14 kPa | 10 | 250 | 100 | 639 | 1 | 10 | 13.694 ± 0.0622 |

**Table S3.** Number of cells in cell area and YAP activity analyses.

|  | **Number of U2OS cells** | **Number of hMSCs** |
| --- | --- | --- |
| **Flat** | 396 | 291 |
| **d1000p5h1** | 48 | 32 |
| **d1000p2.5h1** | 120 | 27 |
| **d500p5h1** | 28 | 42 |
| **d500p2.5h1** | 58 | 27 |
| **d500p1h1** | 120 | 22 |
| **d200p5h1** | 38 | 34 |
| **d200p2.5h1** | 89 | 29 |
| **d200p1h1** | 59 | 25 |
| **d1000p5h3** | 58 | 46 |
| **d1000p2.5h3** | 45 | 36 |
| **d500p5h3** | 46 | 15 |
| **d500p2.5h3** | 58 | 86 |
| **d500p1h3** | 31 | 15 |
| **d200p5h3** | 59 | 24 |
| **d200p2.5h3** | 69 | 34 |
| **d200p1h3** | 60 | 51 |
| **14 kPa** | 134 | 63 |
| **7 kPa** | 164 | 62 |
| **3 kPa** | 86 | 84 |
| **1 kPa** | 23 | 63 |

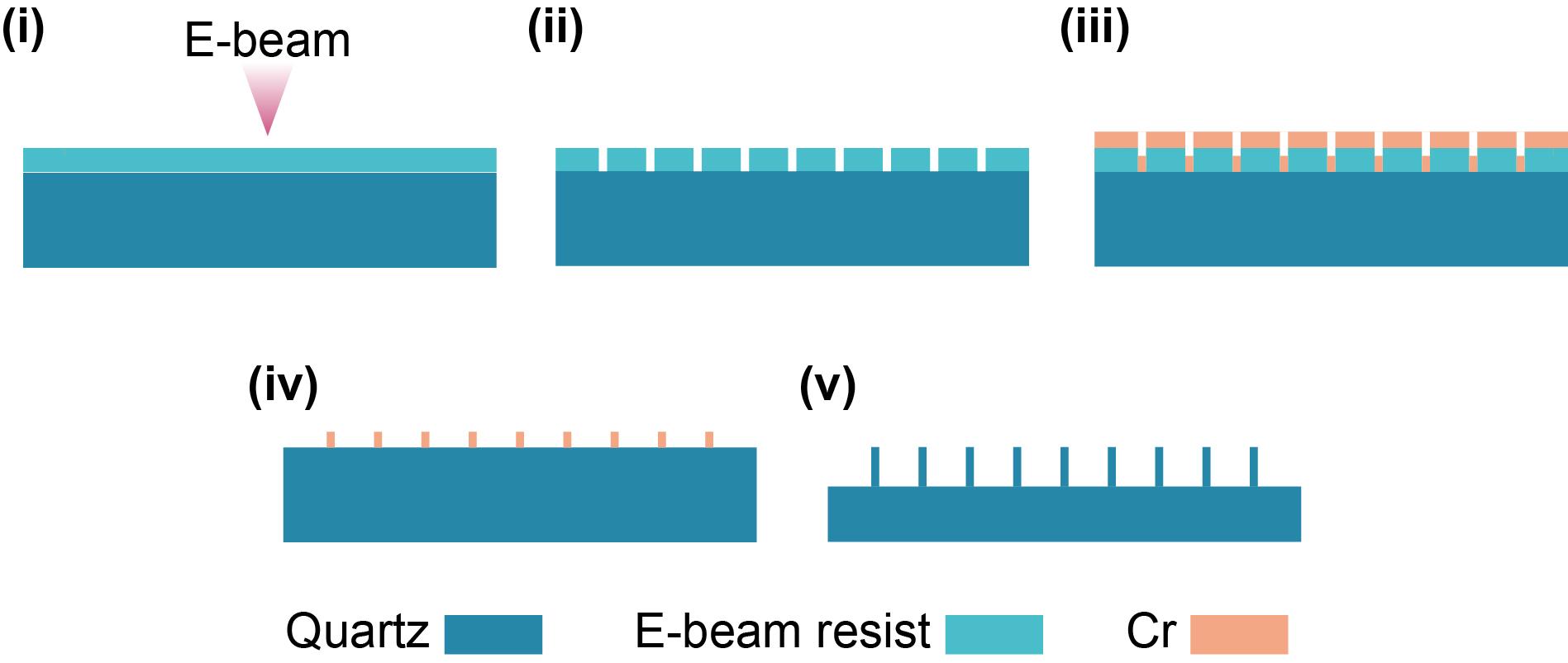

**Figure S1.** Schematics of the nanofabrication process. (i) After the surface of a quartz chip was cleaned and subsequently coated with electron beam (E-beam) resist, designed patterns were written on the chip surface with a JOEL E-beam system. (ii) After development, the exposed E-beam resist was removed, leaving patterns on the chip surface. (iii) A layer of chromium (Cr) was deposited on the chip surface. (iv) Part of the Cr layer was removed via a lift-off process, leaving Cr patterns on the chip surface. (v) The chip was dry-etched with the Cr patterns as masks, and the Cr patterns were removed with an etchant after dry-etching.

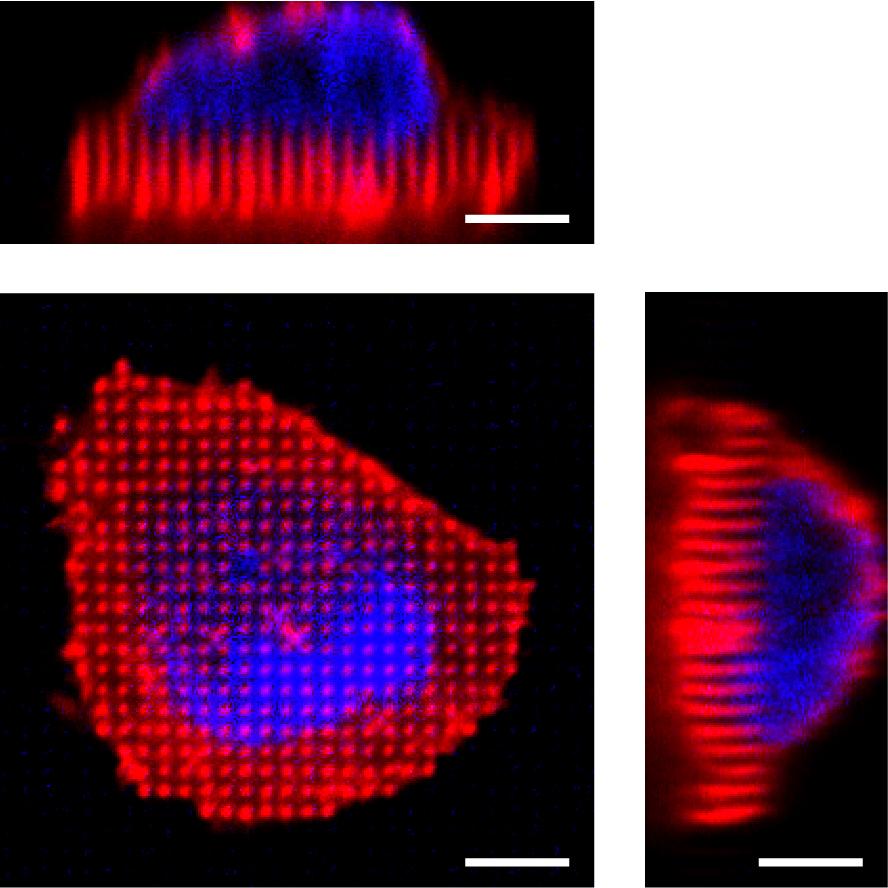

**Figure S2.** Confocal fluorescence images of U2OS cells expressing a membrane marker RFP-CAAX cultured on d200p1h3 nanopillars. The top and right-side images are the corresponding x-z and y-z views showing the deformation of the cell membrane around each nanopillar. The cell was also stained with Hoechst to indicate the nucleus (blue). Scale bars: 5 μm.

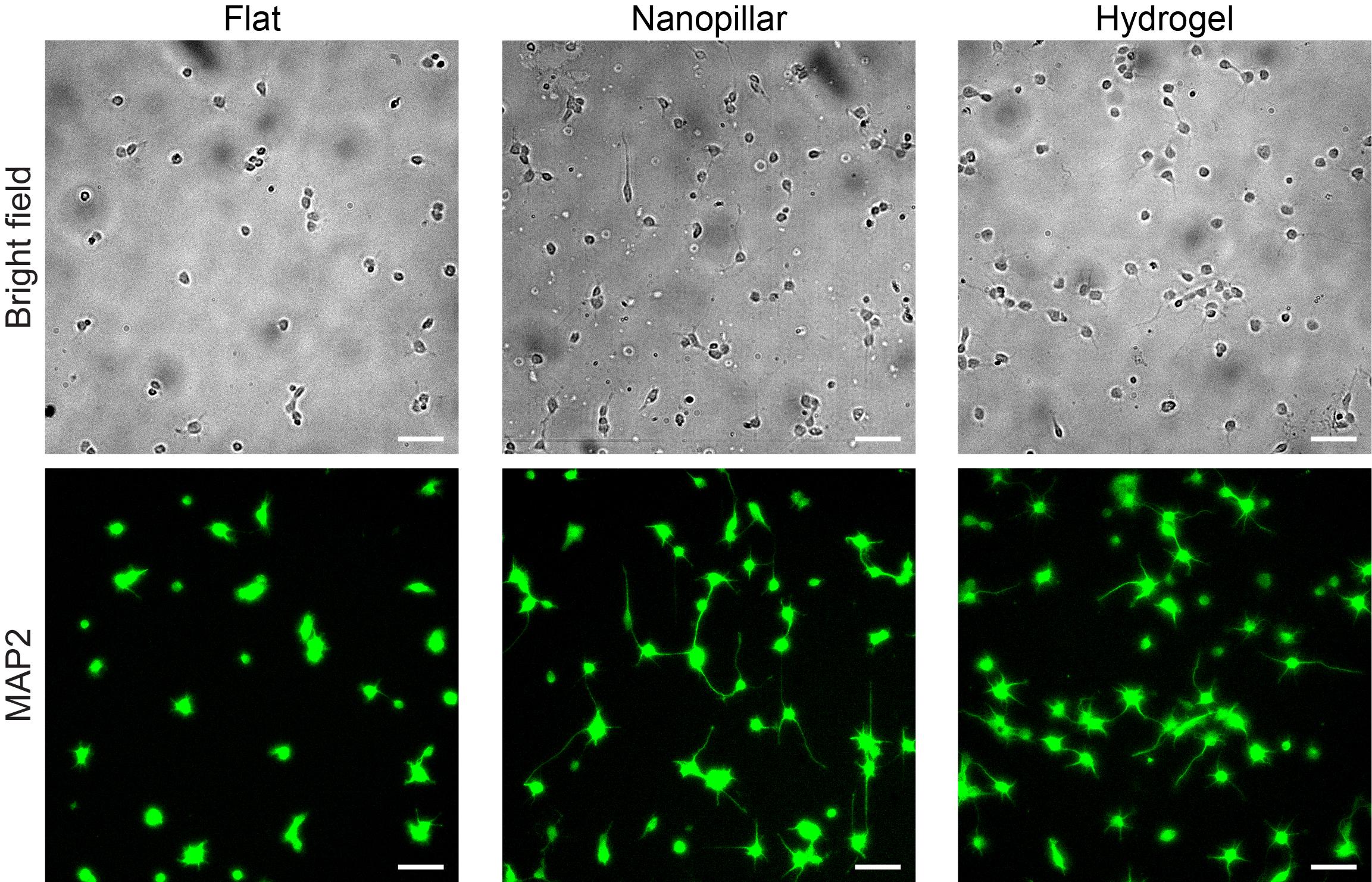

**Figure S3.** Large field-of-view images of hippocampal neurons on flat quartz, nanopillar quartz, and hydrogel surfaces. E18 rat hippocampal neurons were stained with anti-Map2 after 20 hr of culture on flat quartz (flat), nanopillar quartz (nanopillar), and 1 kPa hydrogel (hydrogel) surfaces. Scale bars: 50 μm.

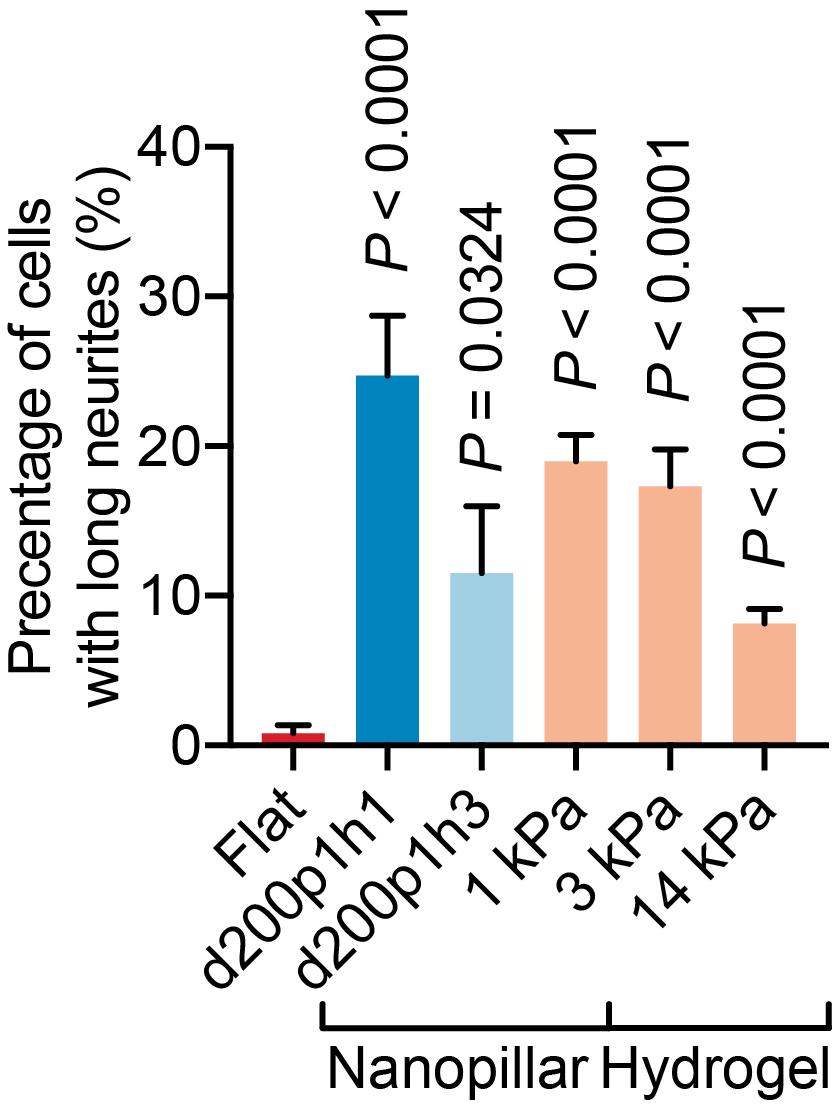

**Figure S4.** Neurite outgrowth on varied substrates. Percentages of hippocampal neurons with at least one neurite longer than 35 μm (n = 193, 297, 95, 178, 195, 255 cells in flat, d200p1h1, d200p1h3, 1 kPa, 3 kPa, and 14 kPa groups, respectively; mean ± s.e.m.) were quantified. *P* value in comparison to flat surface was determined by unpaired two-tailed t test.

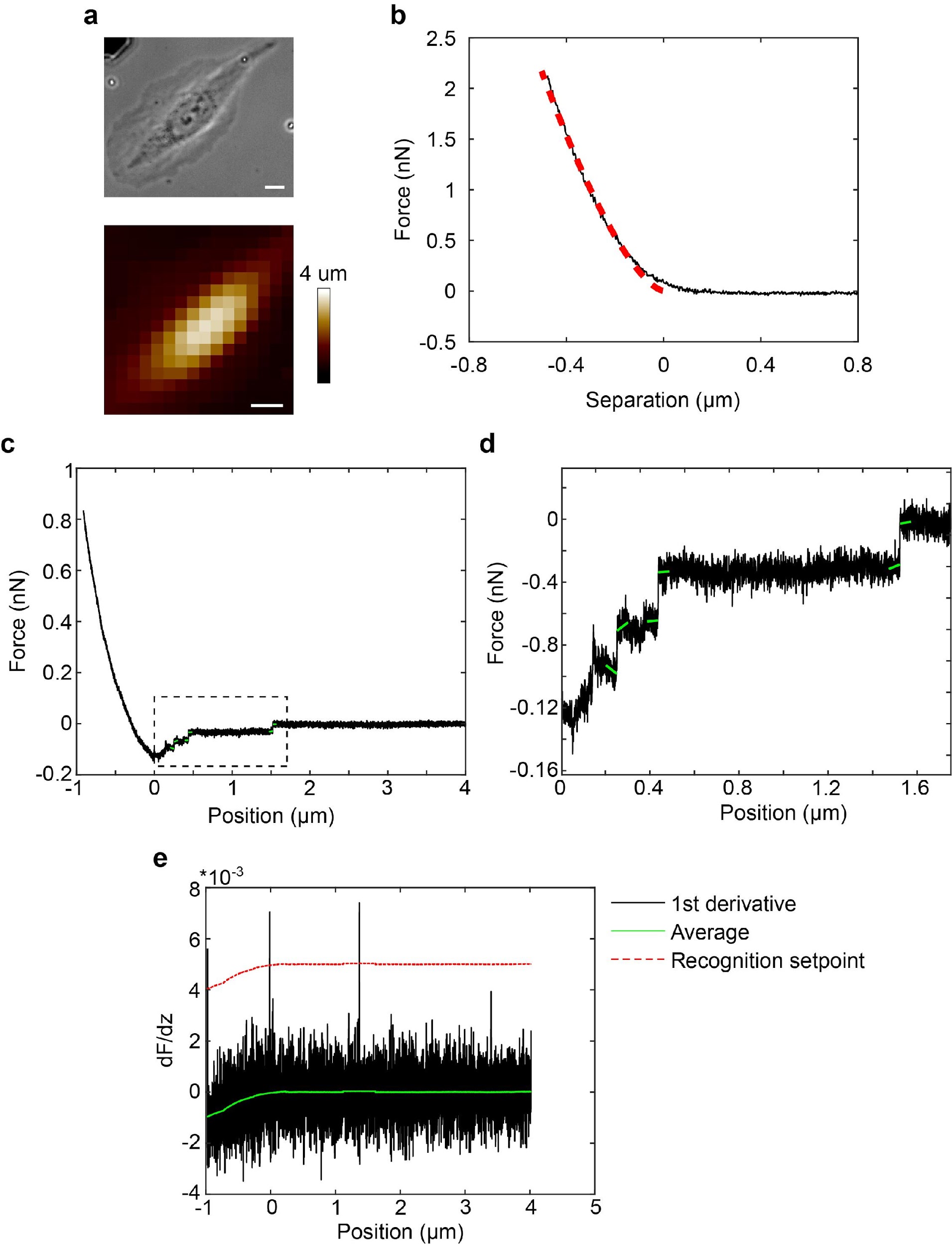

**Figure S5.** AFM measurements. (a) An optical image (upper) and an AFM-obtained topography map (lower) of a U2OS cell on a flat quartz surface. (b) A force-separation curve (black line) with a Hertz indentation model fit (dashed red line) of AFM indentation on a U2OS cell grown on flat quartz surface. Fig. 3b and Extended Data Fig. 5a have a similar area, but their force-separation curves show a clearly lower cell stiffness on nanopillars. © A representative retraction curve showing multiple membrane tether events. (d) Zoom-in view of the tether rupture events. Tether rupture events occur after 250, 450, and 1500 nm. (e) Derivative of the force curve used to identify tether rupture events. Scale bars: 10 μm (a).

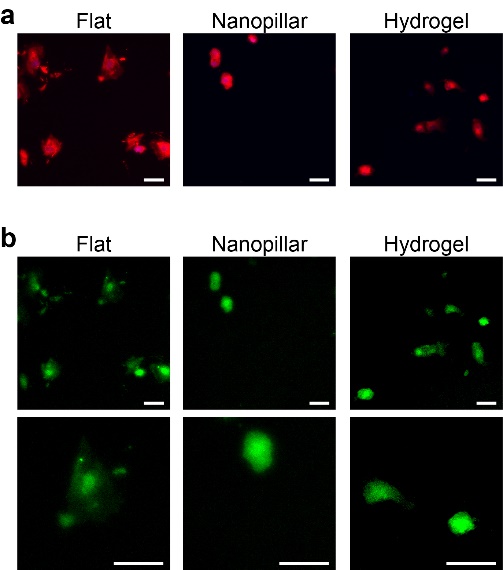

**Figure S6.** Quartz nanotopography reduces cell area and YAP nuclear localization of hMSCs in a manner similar to soft hydrogels. (a) Fluorescence images of hMSCs with CellMask (red for cell membrane) staining on flat quartz, nanopillar quartz, and 1 kPa hydrogel surfaces. (b) Fluorescence images of hMSCs with anti-YAP antibody staining images (top row) and zoomed-in images (bottom row) on flat quartz, nanopillar quartz, and 1 kPa hydrogel surfaces. Lower images show the zoom-in views of the upper images. Scale bars: 50 μm (a, b).

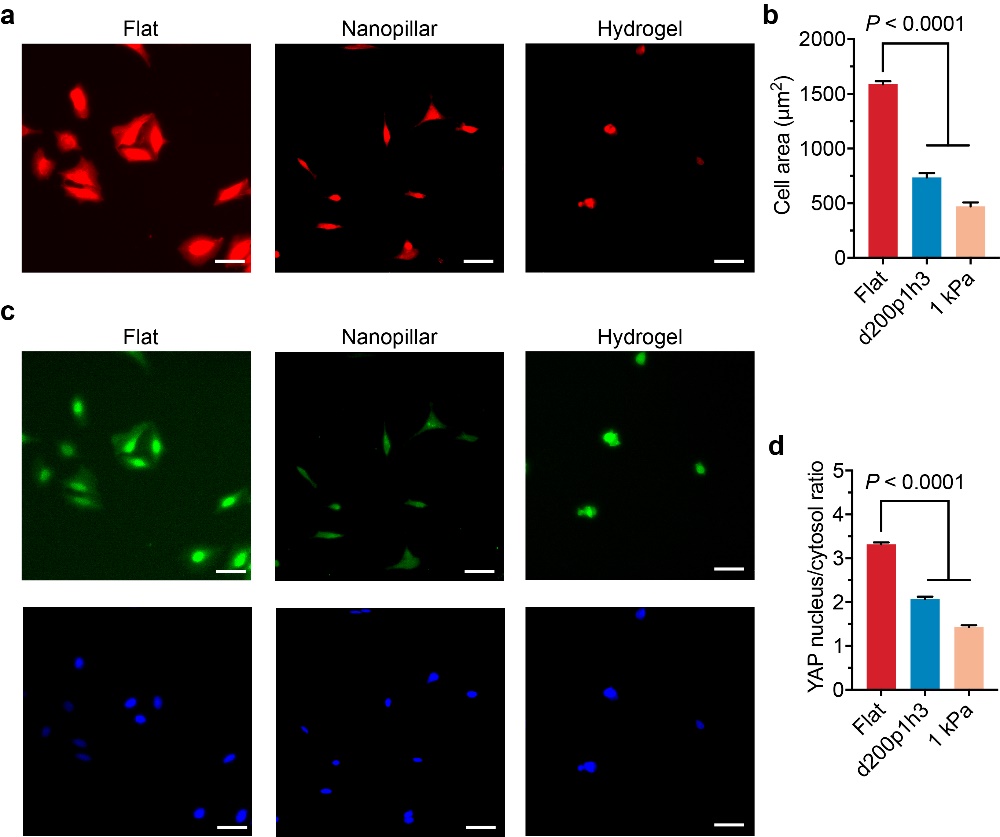

**Figure S7.** Quartz nanotopography reduces cell area and YAP nuclear localization of U2OS in a manner similar to soft hydrogels. (a) Fluorescence images of U2OS cells with CellMask (red for cell membrane) staining on flat quartz, nanopillar quartz, and 1 kPa hydrogel surfaces. (b) Quantitative analysis of cell area on flat quartz, nanopillar quartz, and 1 kPa hydrogel surfaces. (c) Fluorescence images of U2OS cells with anti-YAP antibody (green for YAP) and Hoechst (blue for nuclei) staining. (d) Quantitative analysis of YAP nucleus/cytosol ratio on flat quartz, nanopillar quartz, and 1 kPa hydrogel surfaces. (n= 396, 60, 23 cells in flat, nanopillar, and hydrogel groups, respectively; mean ± s.e.m.). *P* value in comparison to flat surface was determined by unpaired two-tailed t test (b, d). Scale bars: 50 μm (a, c).

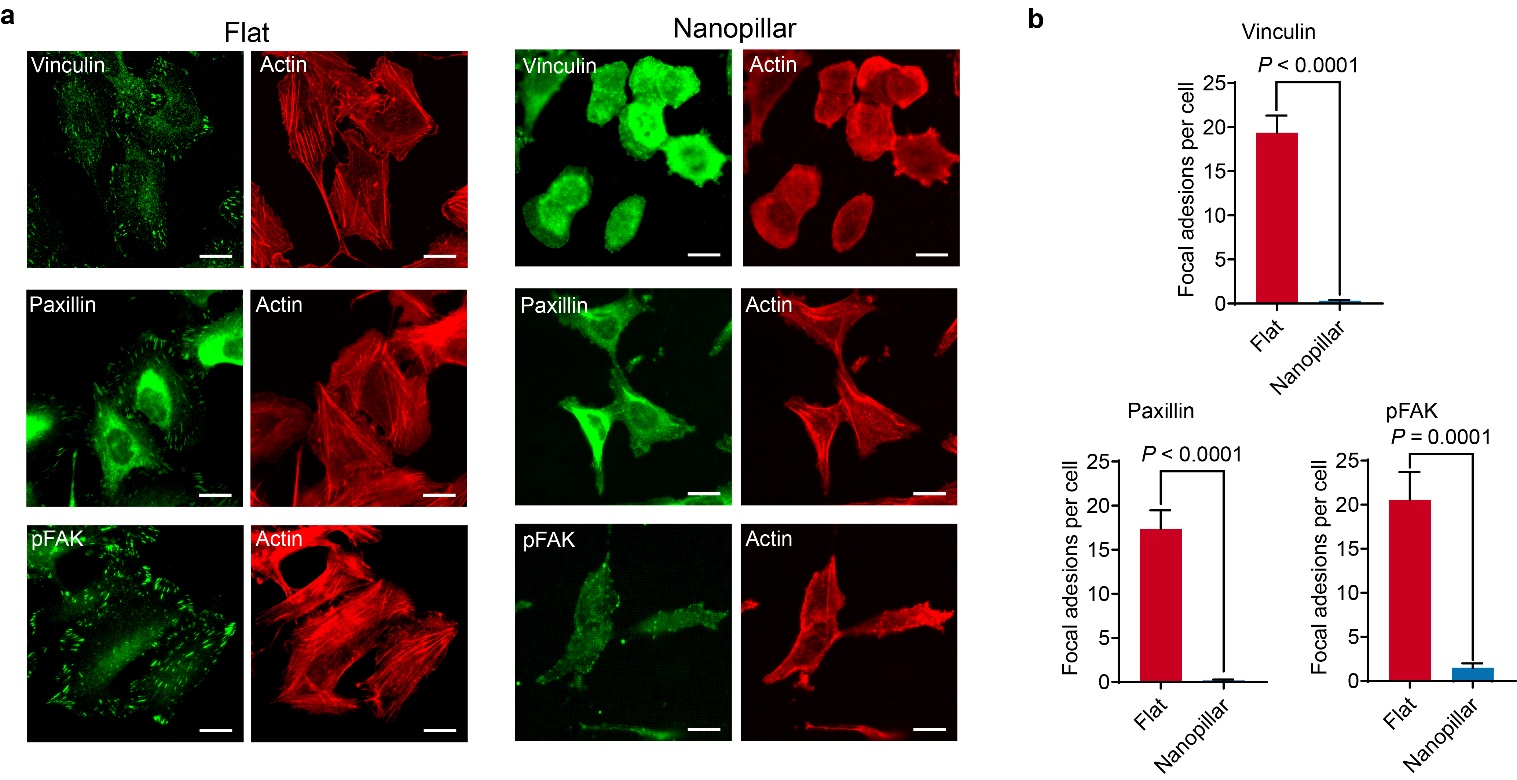

**Figure S8.** Nanotopography reduces focal adhesions and stress fibers. (a) Antibody staining of focal adhesion proteins including vinculin, paxillin, and pFAK along with actin in U2OS cells. Actin was stained with phalloidin dye (red). Focal adhesion proteins were stained with corresponding antibodies and fluorescently labeled (green). (b) Quantitative analysis of focal adhesion patches (larger than 1 μm^2^) on flat quartz, nanopillar quartz, and 1 kPa hydrogel surfaces (n = 34, 51, and 18 cells in flat, nanopillar, and hydrogel groups, respectively; mean ± s.e.m.). Scale bars: 20 μm.

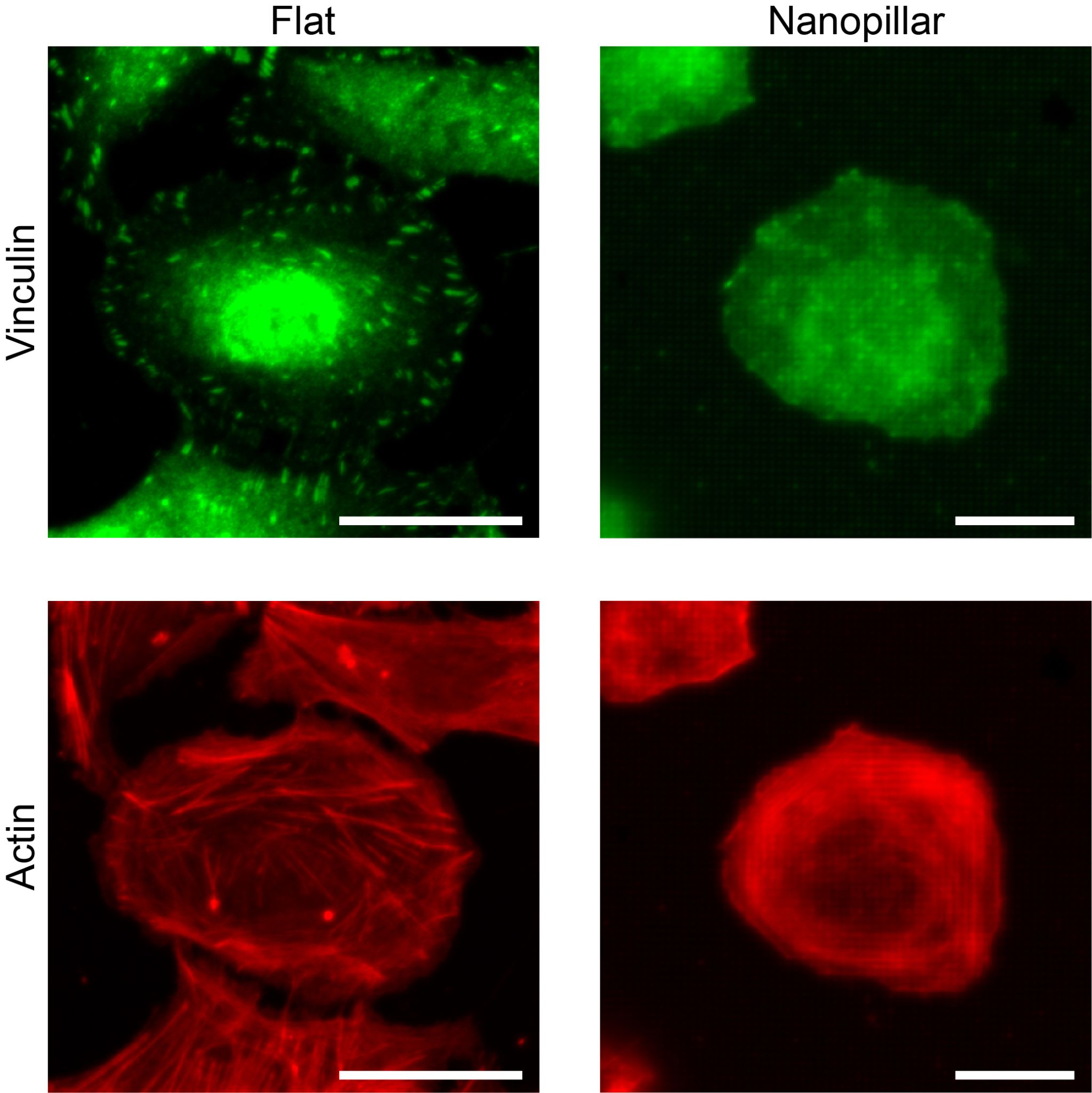

**Figure S9.** Confocal imaging of Z-stack of stress fibers and vinculin. The image stack was flattened to show that the reduced stress fibers and focal adhesions (by vinculin staining) were not due to limited depth of fluorescence imaging.

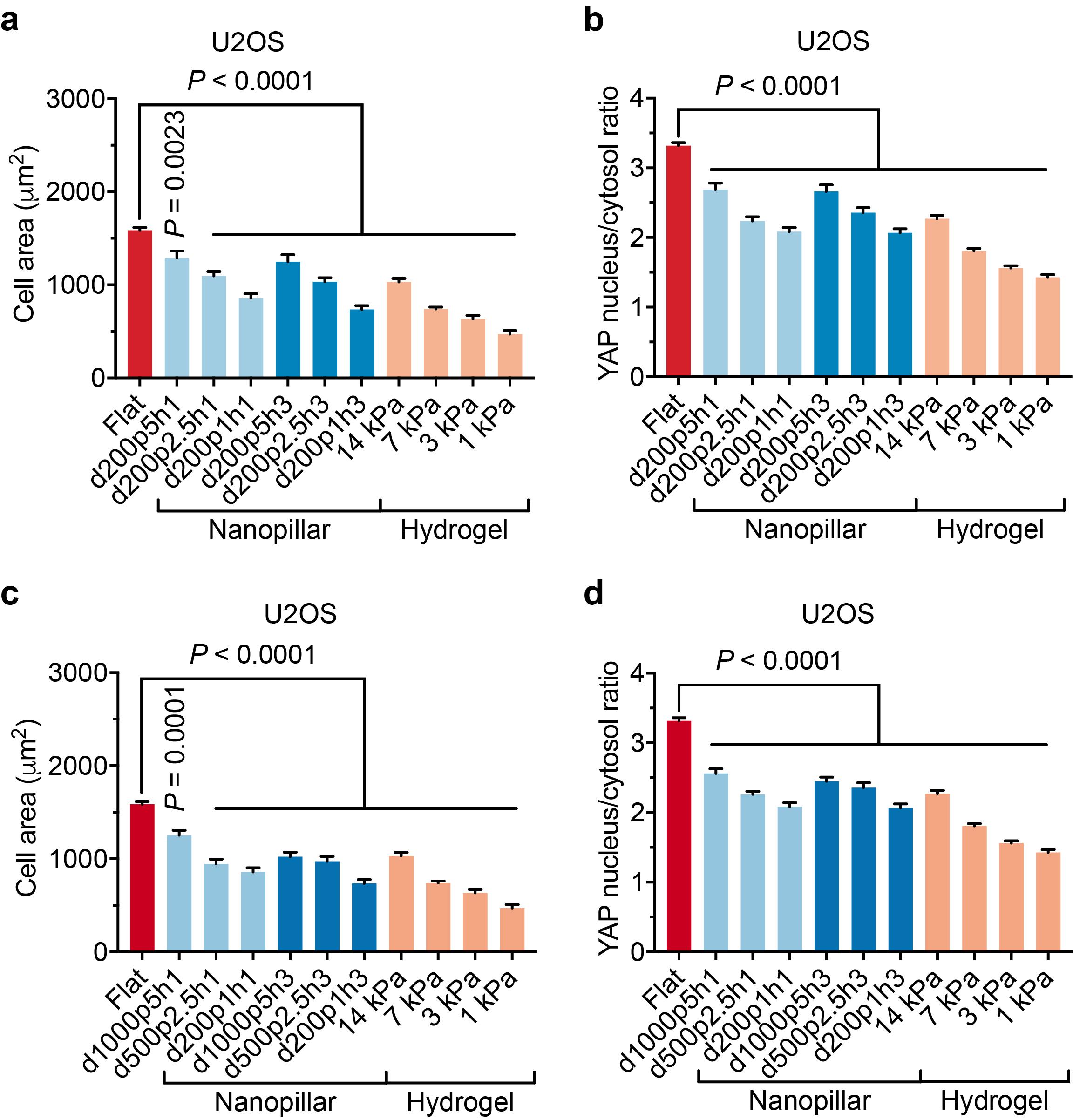

**Figure S10.** Cell behavior can be adjusted with nanopillar dimensions. (a, b) Quantitative analyses of cell area (a) and YAP nucleus/cytosol ratio (b) of U2OS cells on 200-nm-diameter nanopillars with varied pitches and heights. (n=23 to 396 cells, Table S3; mean ± s.e.m.). (c, d) Quantitative analyses of cell area (c) and YAP nucleus/cytosol ratio (d) of U2OS cells on nanopillars with varied diameters and heights but a constant diameter/pitch ratio. (n=23 to 396 cells, Table S3; mean ± s.e.m.). *P* value in comparison to flat surface was determined by unpaired two-tailed t test.
